# A human prenatal skin cell atlas reveals immune cell regulation of skin morphogenesis

**DOI:** 10.1101/2023.10.12.556307

**Authors:** Nusayhah Hudaa Gopee, Ni Huang, Bayanne Olabi, Chloe Admane, Rachel A. Botting, April Rose Foster, Fereshteh Torabi, Elena Winheim, Dinithi Sumanaweera, Issac Goh, Mohi Miah, Emily Stephenson, Win Min Tun, Pejvak Moghimi, Ben Rumney, Peng He, Sid Lawrence, Kenny Roberts, Keval Sidhpura, Justin Englebert, Laura Jardine, Gary Reynolds, Antony Rose, Clarisse Ganier, Vicky Rowe, Sophie Pritchard, Ilaria Mulas, James Fletcher, Dorin-Mirel Popescu, Elizabeth Poyner, Anna Dubois, Andrew Filby, Steven Lisgo, Roger A. Barker, Jong-Eun Park, Roser Vento-Tormo, Phuong Ahn Le, Sara Serdy, Jin Kim, CiCi Deakin, Jiyoon Lee, Marina Nikolova, Neil Rajan, Stephane Ballereau, Tong Li, Josh Moore, David Horsfall, Daniela Basurto Lozada, Edel A. O’Toole, Barbara Treutlein, Omer Bayraktar, Maria Kasper, Pavel Mazin, Laure Gambardella, Karl Koehler, Sarah A. Teichmann, Muzlifah Haniffa

## Abstract

Human prenatal skin is populated by innate immune cells including macrophages, and whether they act solely in immunity or have additional functions in morphogenesis is unclear. We assembled the first comprehensive multi-omic reference atlas of prenatal human skin (7-16 post-conception weeks), combining single cell and spatial transcriptomic data, to characterise the skin’s microenvironmental cellular organisation. This revealed that crosstalk between non-immune and immune cells underpins formation of hair follicles, has implications for scarless wound healing, and is critical for skin angiogenesis. We benchmarked a skin organoid model, derived from human embryonic stem (ES) and induced pluripotent stem (iPS) cells, against prenatal and adult skin, demonstrating close recapitulation of the epidermal and dermal skin components during hair follicle development. Notably, the skin organoid lacked immune cells and had markedly diminished endothelial cell heterogeneity and quantity. From our *in vivo* skin cell atlas data, we found that macrophages and macrophage-derived growth factors play a key role in driving endothelial development prenatally. Indeed, vascular network formation was enhanced following transfer of autologous iPS-derived macrophages into both endothelial cell angiogenesis assays and skin organoid cultures. In summary, innate immune cells moonlight as key players in skin morphogenesis beyond their conventional immune roles, a function they achieve via extensive crosstalk with non-immune cells. Finally, we leveraged our human prenatal skin cell atlas to further our understanding of the pathogenesis of genetic hair and skin disorders.

## Introduction

Human skin organogenesis begins after gastrulation from two primary germ layers. The epidermis, the most superficial layer of the skin, melanocytes and neuronal cells arise from ectodermal differentiation. The dermis, which lies just deep to the epidermis and is separated by the basement membrane, and the endothelial and mural cells it contains, differentiate from the embryonic mesoderm across most anatomical sites, except in facial and cranial skin where dermal cells arise from ectoderm-derived neural crest cells^1,2^. The epidermis is initially composed of a single layer of ectodermal cells^3^. By 4 post conception weeks (PCW), two layers can be observed: a basal cell layer and an outer layer known as the periderm that represents the first permeability barrier^4^. Cells from the periderm are shed into the amniotic fluid during the second trimester when the basal layer begins stratification^5^.

The skin appendages, which include hair follicles and sebaceous glands, form in a cephalo-caudal direction during prenatal life^6,7^. Hair follicle morphogenesis is initiated by the interaction between epidermal placodes (focal sites of epidermal layer thickening) and dermal condensates (aggregates of dermal fibroblasts). With these interactions, the prenatal hair follicle develops from the epidermal placode that penetrates the dermis around 11-14 PCW^4,6^. Subsequently, the keratinised hair shaft forms within the centre of the hair follicle, which is surrounded by an epidermal hair sheath and a dermal root sheath^1^. Sebaceous glands, the sebum secreting glands of the hair follicle, start forming from around 16 PCW, and prenatal hair that protrudes from the skin is observed around 18 PCW^6^. There is, however, a paucity of information about the precise cellular composition of human prenatal skin over these developmental periods and whether cells interact in functional microanatomical niches that support skin morphogenesis.

The skin performs key immune functions as a barrier organ after birth. Unlike postnatal skin that is exposed to air and a microbial rich environment, prenatal skin interfaces with the amniotic fluid in a sterile environment^8^. However, immune cells such as macrophages seed the skin as early as 6 PCW^5^ and express a range of pro-inflammatory genes, although the expression of major histocompatibility complex class II (MHC-II) genes, relating to antigen presentation, is only upregulated after 11 PCW^9^. Macrophages are elsewhere known to regulate various aspects of tissue homeostasis, contributing to angiogenesis in disease setting^8,10^, murine hair follicle development and cycling^11^ and cutaneous wound repair^5,12^. The decoupling of expression of pro-inflammatory genes from MHC-II genes before 11 PCW^9^ suggests that antigen presentation may not be a key function of human macrophages during early gestation. Together with evidence for their role in tissue homeostasis and healing in murine models, this raises the question as to whether macrophages in fact contribute to human early skin morphogenesis.

Our study is the first comprehensive multi-omic cell atlas of 7-16 PCW human prenatal skin. We profiled human prenatal skin using single-cell RNA sequencing (scRNA-seq), spatial transcriptomics and multiplex RNA *in situ* hybridisation to decode the dynamic cellular and molecular changes across gestation that regulate human skin and hair follicle morphogenesis. We leveraged adult healthy skin and hair follicle datasets^13,14^ for comparison with prenatal skin to assess developmental specific features contributing to scarless skin healing and cues guiding *de novo* hair follicle formation. We also performed comparative analyses with a hair-bearing skin organoid model^15^ to establish the faithfulness of skin organoids in recapitulating human skin development and identify molecular mechanisms that can further enhance skin organoid models in future experimental settings. We uncovered an important role for macrophages in prenatal skin vascular network formation, which we functionally validated *in vitro*.

## Results

### A single cell atlas of human prenatal skin

To characterise the role of distinct lineages and cell states in the development of human prenatal skin, we obtained single cell suspensions of skin from 7 to 16 PCW, spanning the first and second trimesters, during which skin architecture matures and hair follicles first develop (**Fig. 1a**). Prenatal skin cells were isolated by fluorescence-activated cell sorting (FACS) into CD45^+^ and CD45^-^ fractions, allowing the selection of live, single immune and non-immune populations respectively, for scRNA-seq profiling (10X Genomics) (**Extended Data Fig. 1a, Supplementary Table 1**). In addition, to enhance keratinocyte and endothelial cell capture, we isolated all cells that were not within the CD34^+^CD14^-^ fraction from two prenatal skin samples (**Extended Data Fig. 1a, Supplementary Table 1)**. Single-cell T-cell receptor alpha and beta sequencing (abTCR-seq) data was also generated to accurately resolve T-cell subsets. Spatial validation was carried out using multiplex RNA *in situ* hybridisation (RNAScope), newly generated spatial transcriptomic (Visium) data from embryonic facial and abdominal skin, and published Visium data from embryonic limb from which only skin areas were analysed^16^ (**Fig. 1a**). In addition, we integrated new and published single-cell datasets of a hair-bearing skin organoid model, derived from human embryonic stem (ES) and induced pluripotent stem (iPS) cells^15^, and adult skin^13^ for comparative analysis (**Fig. 1a**). We also compared *in vivo* prenatal and organoid hair follicle cells with scRNA-seq data of adult hair follicles^14^. Our data can be explored interactively through our web portal which leverages WebAtlas^17^ for intuitive visualisation and query (cell type and gene expression) of our single-cell, spatial and integrated datasets (https://developmental.cellatlas.io/fetal-skin; password: fs2023). The analysis software for this study is archived at Zenodo (https://doi.org/10.5281/zenodo.8164271).

**Fig. 1:**
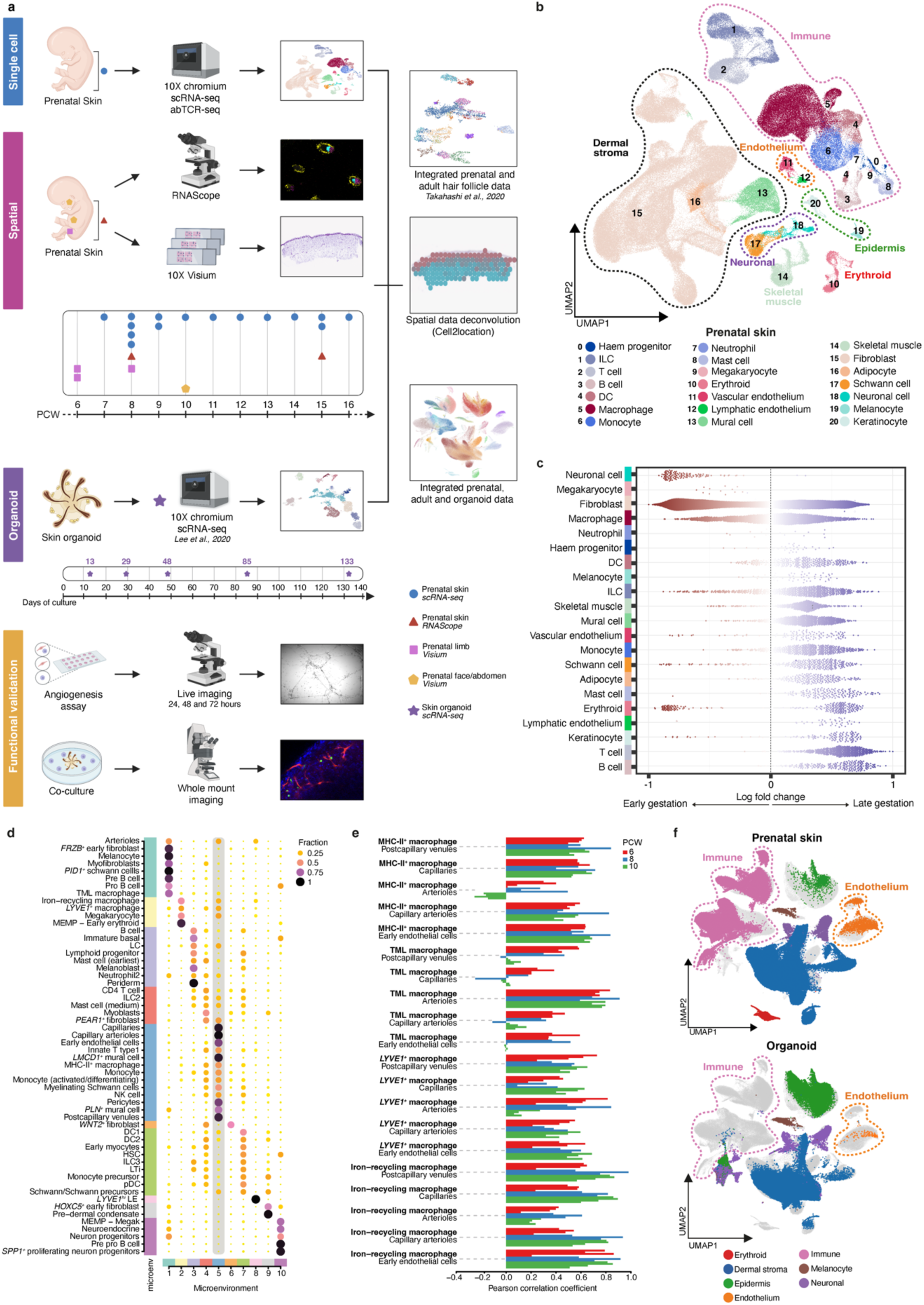
A single cell atlas of human prenatal skin. **(a)** Experimental overview demonstrating the generation of scRNA-seq data from dissociated prenatal skin cells (n=15, 7-16 PCW). Spatial experiments were carried out using RNAScope and Visium, and a hair-bearing skin organoid dataset was integrated for benchmarking. Findings of the study were functionally validated using an angiogenesis assay and skin organoid co-culture. **(b)** UMAP visualisation of the prenatal skin dataset with broad annotation of cell states, as denoted by colour and number in the legend. **(c)** Milo beeswarm plot showing differential abundance of neighbourhoods in prenatal skin across gestation time, annotated by broad cell labels. Red/blue neighbourhoods are significantly enriched in earlier/later gestation respectively. Colour intensity denotes degree of significance. **(d)** Dotplot showing spatial microenvironments. Cell type to microenvironment coefficients are normalised by cell type sums; cell type to microenvironment assignment is shown by colour. Microenvironment 5, which shows co-locating macrophages and endothelial cells, is highlighted. **(e)** Bar plot showing cell type co-location, indicated by positive Pearson correlation coefficient, for selected cell type pairs (macrophage and endothelial cells). Pearson correlation coefficients were calculated across all skin-covered spots of Visium samples; each sample is shown by an individual bar. **(f)** UMAP visualisations of the integrated prenatal skin, adult skin^13^ and skin organoid^15^ datasets, coloured by broad cell lineages. ASDC: *Axl+Siglec6+* dendritic cells; DC: dendritic cells; HSC: hematopoietic stem cells, ILC: innate lymphoid cells, LC: Langerhans cells, LTi: lymphoid tissue inducer cells, pDC: plasmacytoid dendritic cells, TML macrophage: *TREM2^+^* microglia-like macrophage.

Our prenatal skin scRNA-seq dataset comprised 235,201 cells of which 186,582 cells passed quality control and doublet exclusion (**Extended Data Fig. 1b**). Maternal cell contamination was removed using Souporcell (49 cells). Broad cell labels (epidermis, dermal stroma, immune and endothelium) were assigned following dimensionality reduction analysis and batch correction (see Methods) (**Fig. 1b, Extended Data Fig. 1c**). Further dimensionality reduction and clustering was performed on the broad cell clusters prior to fine-grained manual annotation of cell states (**Supplementary Table 2**). Graph-based differential abundance analysis using Milo^18^ was performed to test changes in cellular composition across gestation. The enrichment of different cell populations varied with gestational stages. Amongst ectoderm-derived cells, neuronal cells and periderm were enriched in early gestation whilst suprabasal and hair follicle epidermal cells were mainly observed in later gestation (**Fig. 1c, Extended Data Fig. 1d, Supplementary Table 3**). Mesoderm-derived cells, including skin fibroblasts and endothelial cells, and immune cells were present throughout gestation (**Fig. 1c, Extended Data Fig. 1d, Supplementary Table 3**). Innate immune cells, such as macrophages and innate lymphoid cells (ILCs), were present from early gestation whilst B- and T-cells emerged later, accompanying thymus, bone marrow and spleen formation from around 10 PCW (**Fig. 1c, Extended Data Fig. 1d, Supplementary Table 3**). Some subsets of macrophages, innate lymphoid cells (ILCs) and fibroblasts, exhibited distinct gene expression profiles between early and late gestation, suggesting functional evolution during development or dual waves of production (**Fig. 1c, Extended Data Fig. 1d, Supplementary Table 3**).

To locate cells identified from scRNA-seq *in situ,* we performed Cell2location^19^ analysis using spatial transcriptomic data from facial and abdominal skin (10 PCW) and embryonic lower limb skin (6-8 PCW)^16^. We assessed if specific cell types were more likely to be co-located within microanatomical tissue niches (microenvironments) using non-negative matrix factorisation and correlation analyses (see methods). Co-location was indicated by a high proportion of two or more cell types sharing a microenvironment (**Fig. 1d**) and/or by positive correlation coefficient between cell pairs (**Fig. 1e, Extended Data Fig. 1e**). Indeed, we found that prenatal skin was made up of several microenvironments comprising epidermal, dermal, vascular, and neuronal cells (**Fig. 1d, e, Extended Data Fig. 1e**). Each of these microenvironments included specific types of immune cells (**Fig. 1d**). For example, pre-dermal condensate (pre-Dc) co-located with dendritic cells and lymphoid cells in an ‘early hair follicle microenvironment’, while macrophages co-located with endothelial and neuronal cells in a ‘neurovascular microenvironment’ (**Fig. 1d, e, Extended Data Fig. 1e**). These observations led us to question whether immune cells occupy distinct microanatomical niches where they undertake a moonlighting function distinct from their role in immunity during early development. We hypothesised a role for immune cells in supporting human skin morphogenesis and specifically macrophages in skin vascular network formation, akin to the role of zebrafish macrophages in guiding haematopoietic stem and progenitor cells in vascular niches^10^.

We next integrated and compared our human prenatal and adult skin data^13^ with the skin organoid model^15^ to determine faithfulness of the organoid model to *in vivo* skin and its potential utility to functionally assess the role of immune cells in skin morphogenesis (**Extended Data Fig. 2a, b**). Using the same approach as for prenatal skin, the skin organoid data was analysed and assigned broad (epidermis, dermal stroma and endothelium) and refined cell annotations. Skin organoid had the same ectoderm-derived cell types as observed in prenatal skin with basal cells emerging prior to suprabasal cells (**Fig. 1f, Extended Data Fig. 2a, b**). However, skin organoid had a more restricted mesoderm-derived cellular repertoire with a paucity of endothelial cells and absent immune cells, in keeping with most iPS-derived peripheral tissue organoid models (**Fig. 1f, Extended Data Fig. 2a, b**).

To determine to what extent the skin organoid recapitulates human skin differentiation at a molecular level, we assessed similarity between prenatal skin, adult skin, and skin organoid for broad cell categories, based on transformed gene expression levels and logistic regression analyses (see methods) (**Extended Data Fig. 2c, d, Supplementary Table 4**). Broadly, cell states were conserved between skin organoid, prenatal and adult skin, but skin organoid cell states matched prenatal skin more closely than adult skin by gene expression profile across culture duration (**Extended Data Fig. 2c, d, Supplementary Table 4**). However, the tempo of differentiation varied across the skin cell lineages. 12-week-old organoid fibroblasts most closely aligned to 7-8 PCW skin fibroblasts, and even after 19 weeks of culture, fibroblasts, mural and Schwann cells had a low probability of correspondence to adult skin cell states (**Extended Data Fig. 2d, Supplementary Table 4**). In contrast, accelerated differentiation was observed in keratinocytes and melanocytes, with alignment to adult cell states seen as early as 4 weeks of organoid culture (**Extended Data Fig. 2d, Supplementary Table 4**). The ES/iPS-derived skin organoid model recapitulated the different components of prenatal skin hair follicle, interfollicular epidermis, neuronal cells, and dermal fibroblasts but immune cells were not represented, and endothelial cells were markedly diminished.

### Hair follicle - epidermal placode and matrix formation

Hair follicle formation is initiated by interactions between dermal (mesenchymal) and epidermal (epithelial) compartments which result in the formation of the epidermal-derived hair placode and matrix and in the differentiation of hair-specialised fibroblasts^20,21^. The precise mechanisms of *de novo* hair follicle formation in human embryonic development are poorly understood and are largely inferred from murine studies^22^. Human studies have mainly been limited to morphological descriptions during development^23–26^ or focused on adult skin where established hair follicles cycle through the distinct phases of hair growth (anagen), regression (catagen) and resting (telogen)^27,28^. Our single cell dataset captures the onset of hair follicle formation enabling direct comparison between prenatal developing and adult cycling hair follicles.

Prenatal skin up to 8 PCW consisted of a layer of epidermal cells overlying the dermal stroma and an outer epithelial periderm was observed to slough from the prenatal skin as early as 11 PCW (**Fig. 2a)**. At 14-15 PCW, budding of basal cells (hair placodes/germs) and elongation of hair follicles (hair pegs) were observed, some of which were seen as circular structures in transverse cross section of the prenatal hair follicles (**Fig. 2a**). Representative skin section at 17 PCW showed longitudinally sectioned, elongated hair follicle structures beneath a stratified epidermal layer (**Fig. 2a**)^7,26^.

**Fig. 2:**
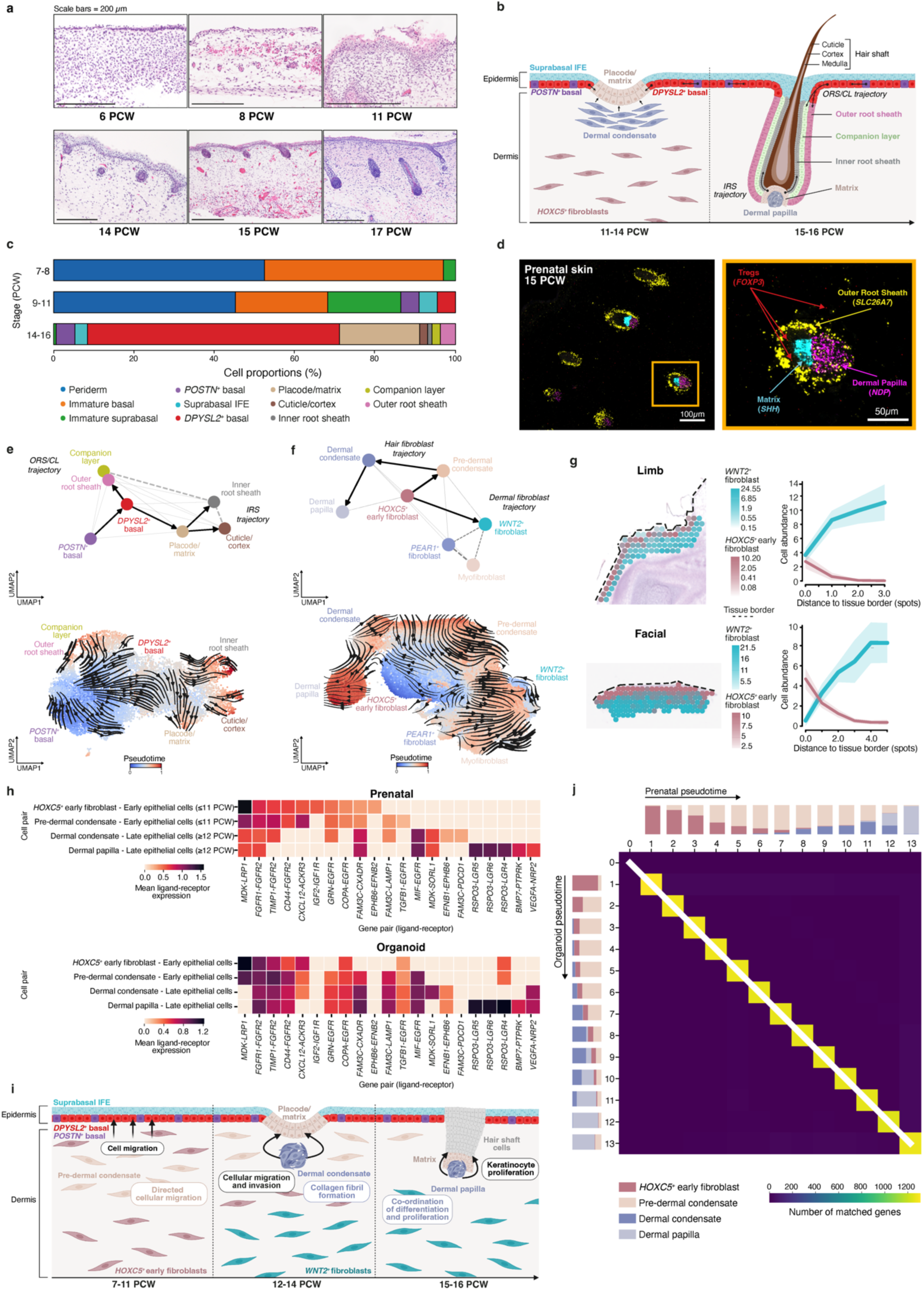
Human prenatal hair follicle development. **(a)** Representative haematoxylin and eosin-stained images showing different developmental stages of prenatal skin and hair follicle morphogenesis. **(b)** Schematic of stages of hair follicle formation. **(c)** Bar plot showing proportions of epidermal cell states across gestational age in prenatal skin. Bar colours represent cell states. **(d)** Large-area (left, scale bar=100μm) and magnified (right, scale bar=50μm) RNAScope images of prenatal skin (representative 15 PCW sample) demonstrating ORS (*SLC26A7*), matrix (*SHH*), dermal papilla (*NDP*) and Tregs (*FOXP3*). **(e)** Inferred pseudotime trajectory of skin organoid epidermal cell states differentiating along the ‘ORS/CL’ and ‘IRS’ trajectories; UMAP overlaid with PAGA and coloured by cell state (top) and overlaid with CellRank state transition matrix inferred arrows and coloured by pseudotime (bottom). **(f)** Inferred pseudotime trajectory of prenatal skin fibroblasts differentiating along the ‘hair’ and ‘dermal’ fibroblast trajectories. UMAP overlaid with PAGA and coloured by cell state (top) and overlaid with CellRank state transition matrix inferred arrows and coloured by pseudotime (bottom). **(g)** Spatial distribution of *WNT2^+^* fibroblasts and *HOXC5^+^* early fibroblasts on two representative prenatal skin Visium samples. Predicted cell abundances shown either as the sum of two colour gradients per spot (left) or averaged across all spots which are located on the same distance from tissue border (right). Shaded areas show 95% confidence interval. **(h)** Heatmap showing significant (adjusted p-value <0.05) CellphoneDB-predicted interactions between prenatal skin hair follicle mesenchymal cells and epithelial cells (early: Immature basal; late: *DPYSL2*^+^ basal, *POSTN*^+^ basal, Placode/matrix, ORS, CL, IRS, Cuticle/cortex). Top 10 interactions per cell pair in prenatal skin are shown (top), with the same interactions plotted from the skin organoid CellphoneDB results (bottom). Colour scale represents the mean expression values of each ligand-receptor pair in corresponding cell pairs. **(i)** Schematic representation of mesenchymal-epithelial signalling and cellular processes during hair formation. **(j)** Aggregate alignment result for all included TFs between *in vitro* skin organoid and *in vivo* prenatal skin (reference) data, shown as a pairwise time point matrix between organoid and reference pseudotime axes. Colour scale represents the number of genes showing a match for the given pair of organoid-reference timepoints; the white line represents the main average alignment path. Stacked bar plots represent the cell compositions at each timepoint (13 equispaced time points on pseudotime [0,1]), coloured by cell types for reference (top) and skin organoid (left). CL: companion layer, IRS: inner root sheath, ORS: outer root sheath.

To delineate early hair follicle development in human prenatal skin, we first sub-clustered and annotated all epidermal and dermal cells (**Extended Data Fig. 2b, Supplementary Table 2**). Consistent with our histological observations, we identified hair follicle cells present from 14 PCW, comprising placode/matrix (*SHH^+^*), outer root sheath (ORS) (*SLC26A7^+^*), companion layer (CL), inner root sheath (IRS), and cuticle/cortex cells (part of the inner layers of the hair follicle) (**Fig. 2a-c, Extended Data Fig. 3a, Supplementary Table 5**). In addition, we observed immature and mature interfollicular epidermal (IFE) cells. Immature IFE cells included periderm, immature basal and immature suprabasal cells, which were present from 7 PCW and disappeared after 11 PCW, during the transition from embryonic to fetal skin (**Fig. 2c**). Mature basal (*POSTN^+^* and *DPYSL2^+^*) and suprabasal IFE cells emerged after 11 PCW (**Fig. 2c, Extended Data Fig. 3a**). Sebaceous gland and apocrine gland cells, which mature after 16 PCW^29,30^, were not captured at these stages. Within the dermal compartment, we observed specialised fibroblasts related to hair follicles, the dermal condensate (Dc) and dermal papilla (Dp), from 12 PCW (**Fig. 2b, Extended Data Fig. 3b**).

Next, we evaluated the epidermal placode, a prenatal-specific cell state that gives rise to the hair matrix and which is absent in established adult hair follicles^31^. Accordingly, the adult hair follicle in growth (anagen) phase^14^ comprised cell states which transcriptionally aligned to the prenatal hair follicle, except for placode cells (**Extended Data Fig. 3c, d**). Adult matrix cells expressed genes associated with mature hair structural components (*KRT85*) and regulation of cell proliferation (*LGALS1*), related to their role in hair shaft formation in the growing hair (**Extended Data Fig. 3e**). Conversely, prenatal and skin organoid placode/matrix cells had a higher expression of genes involved in chemotaxis, such as *CXCL14*, a chemokine previously reported to recruit regulatory T-cells (Tregs)^32^, and in control of autoimmunity (*CD24)*^33–35^, signifying the potential role of Treg accumulation and immune protection in the early stages of placode/matrix differentiation (**Extended Data Fig. 3e**). Tregs have been reported to localise around the hair follicle in late second trimester (23 PCW) and in postnatal skin^36,37^. We identified *FOXP3^+^*Tregs primarily within and around hair follicles, as well as the interfollicular epidermis, from as early as 15 PCW by RNAScope staining (**Fig. 2d**).

To infer the differentiation trajectory of epidermal cells, we performed inferred trajectory and pseudotime analysis (CellRank) using skin organoid data to ensure adequate cell numbers. This predicted the differentiation of *POSTN*^+^ basal epidermal cells into *DPYSL2*^+^ basal cells, prior to bifurcation into the placode/matrix and IRS along one arm (‘IRS trajectory’), and the ORS together with the CL along another arm (‘ORS/CL trajectory’) (**Fig. 2e, Extended Data Fig. 3f, Supplementary Table 6**). Along the ‘ORS/CL trajectory’, we identified new genes upregulated by *DPYSL2^+^* basal cells, such as *SPON2* and *AGR2*, in addition to previously reported genes related to ORS differentiation (*BARX2* and *SOX9)*^21,38,39^ (**Extended Data Fig. 3g, h, Supplementary Table 6**)*. DPYSL2^+^* basal cells which were predicted to differentiate into placode/matrix and IRS, upregulated known matrix markers, *SHH* and *WNT10B*^21,38^, and downregulated *SPON2* and *AGR2* (**Extended Data Fig. 3g, h, Supplementary Table 6**). *SPON2* encodes the integrin ligand mindin and is involved in cellular adhesion and immune response^40,41^. Loss of *AGR2,* a molecular chaperone involved in the assembly of cysteine-rich receptors enriched in hair follicles, has been shown to promote cell migration^42,43^. Our findings suggest that decreased adhesion and increased cellular migration properties in *DPYSL2^+^* basal cells may be involved in matrix specification and dermal invagination.

### Hair follicle - dermal condensate and dermal papilla differentiation

Next, we delineated the dermal cell types that are involved in crosstalk with epidermal cells during early hair follicle development. We captured the mesenchymal cells of the developing hair follicle and identified for the first time the human pre-Dc (**Extended Data Fig. 2b, 3b**), a transitional fibroblast state that, in mice, is involved in embryonic hair follicle formation^44–46^. Pre-Dc cells aggregate to form the Dc, which abuts the placode, and highly expressed *CPZ*, a modulator of the Wnt signalling pathway^47^ (**Supplementary Table 2**). Following invagination of the hair follicle, the Dc becomes encapsulated at its base as the Dp (*NDP*^+^)^21,44^ (**Fig. 2d**). Notably, the pre-Dc and Dc are only present in prenatal skin and not in human and mouse adult skin^44^, with self-renewing dermal cells maintaining the Dp during adult hair follicle cycling^48,49^. These self-renewing dermal cells are located within the dermal sheath, a structure of layered connective tissue surrounding the hair follicle^50^. We did not detect a distinct cluster of dermal sheath cells in 7-16 PCW prenatal skin, although this has been identified in a previous single cell analysis of human fetal scalp from 16 and 17 PCW^51^. This is consistent with the earlier development of hair follicles in the scalp compared to trunk skin^6,31^.

To infer the origin of the pre-Dc, Dc and Dp in human prenatal skin, we performed trajectory analysis (CellRank) of the different fibroblast clusters (**Fig. 2f, Extended Data Fig. 4a, Supplementary Table 6**). *FRZB*^+^ fibroblasts were excluded from this analysis as they are primarily observed in one sample from the earliest gestation stage (7 PCW) (**Extended Data Fig. 3b**). Although rare in prenatal skin, *FRZB*-expressing fibroblasts are present in several other developing organs, including the yolk sac, spleen, and gut (**Extended Data Fig. 4b**). Inferred trajectory analysis predicted that *HOXC5^+^* early fibroblasts (located in the upper dermis (**Fig. 2g**) and absent after 11 PCW (**Extended Data Fig. 3b**)) differentiated along two paths. The first path along ‘hair fibroblast trajectory’ formed hair-specialised fibroblasts (pre-Dc, Dc and Dp) and the second path along ‘dermal fibroblast trajectory’ formed into *WNT2^+^*fibroblasts and *PEAR1^+^* fibroblasts (more abundant after 11 PCW) (**Fig. 2f, Extended Data Fig. 3b, 4a, Supplementary Table 6**).

Analysis of genes differentially expressed along the ‘hair fibroblast’ pseudotime (**Extended Data Fig. 4c, Supplementary Table 6**) provided further insights into the processes involved during fibroblast differentiation into the Dp. Genes involved in regulation of cell adhesion (*ADAMST1*), maintenance of cell-cell contact (*CLDN11*), and directed migration (*CXCL12*) were upregulated as the pre-Dc cells migrated towards the epidermis, suggesting a process of collective migration^52–55^ (**Extended Data Fig. 4c, Supplementary Table 6**). Genes implicated in collagen fibril formation and cell adhesion (*COL6A3, MFAP4, PTK7*) were expressed as the pre-Dc aggregated into the Dc (**Extended Data Fig. 4c, Supplementary Table 6**). Formation of the Dp was characterised by genes such as *RSPO3* and *WNT5A* (**Extended Data Fig. 4c, Supplementary Table 6**). These genes are involved in co-ordinating the differentiation and proliferation of the adjacent hair matrix epithelial cells^56,57^.

We next interrogated the mesenchymal-epithelial interactions that instruct early hair follicle formation. Receptor-ligand analysis using CellPhoneDB predicted an interaction between *CXCL12* (ligand), expressed by pre-Dc cells, with *ACKR3* (receptor) from epidermal basal cells up to 11 PCW (**Fig. 2h, Supplementary Table 7**). *CXCL12* expression by hair follicle dermal cells was accordingly downregulated after 12 PCW (**Extended Data Fig. 4d, e**), suggesting that *CXCL12* interacts with the receptor on epithelial cells to mediate migration of pre-Dc to form the Dc^53,58^. Interestingly, lymphoid tissue inducer (LTi) and Type 3 innate lymphoid (ILC3) cells, which co-located with the pre-Dc (‘early hair follicle microenvironment’) (**Extended Data Fig. 1e**), also appeared to interact with pre-Dc cells via ligand-receptor signals that are implicated in regulation of cellular adhesion and migration (*CXCL12*-*CXCR4/DPP4*)^58,59^ (**Extended Data Fig. 4f, Supplementary Table 7**). This suggests that innate immune cells potentially support pre-Dc migration during early hair follicle development. The Dc, whose formation is accompanied by dermal invagination of the placode/matrix, expressed *FAM3C* and *EFNB1,* which were predicted to interact with *LAMP1*/*CXADR* and *EPHB6* respectively on the placode/matrix, and have been reported to promote cellular migration and invasion^60–62^ (**Fig. 2h, i, Supplementary Table 7**). Finally, *RSPO3* from the Dp was predicted to interact with *LGR4/6* (**Fig. 2h, i, Supplementary Table 7**) in overlying matrix cells (**Fig. 2d**) to contribute to proliferation of hair follicle epidermal cells^56^. Notably, the highlighted interactions were conserved between the mesenchymal and epithelial cells of the skin organoid for corresponding stages during hair follicle formation (**Fig. 2h, Extended Data Fig. 4g, Supplementary Table 7**), providing orthogonal validation of our findings and reinforcing the utility of the skin organoid as an accurate model of prenatal skin development.

We further evaluated the differentiation trajectory alignment between prenatal skin and skin organoid using the Genes2Genes analysis framework^63^ to compare the expression of transcription factors (TFs) along the ‘hair fibroblast’ trajectory (*HOXC5*^+^ early fibroblast to dermal papilla). Alignment was computed at both TF level and cell level (aggregated TF-level alignments) (see methods). Overall, the skin organoid closely recapitulated *in vivo* differentiation of hair follicle dermal cells. We observed an average alignment path of 100% match between pseudotime points at cell level and 95% mean alignment similarity at TF level, signifying strong matching of activated gene regulatory programs between prenatal skin and skin organoid during differentiation from *HOXC5*^+^ early fibroblasts to Dp (**Fig. 2j, Supplementary Table 8**). The limited TFs that were mismatched were attributable to the different origins of dermal cells between prenatal skin and skin organoid. Mammalian patterning homeobox (HOX) genes governing trunk and limb development (*HOXC6*, *HOXA7*, *HOXA10*) were upregulated in prenatal skin^64–66^ (**Extended Data Fig. 4h, i, Supplementary Table 8**). In contrast, neuronal and craniofacial development genes (*SIX2*, *LHX6*, *POU3F3*, *HMX1*) were upregulated in skin organoids where dermal cells were derived from cranial neural crest differentiation^15^ (**Extended Data Fig. 4h, i, Supplementary Table 8**).

We also assessed the expression profiles of genes previously reported in mouse hair follicle formation in human prenatal hair follicle epidermal and dermal cells. We identified upregulation of similar signalling pathways, including Wnt/Eda in the initial stages of hair placode specification, fibroblast growth factor 20 (*FGF20*) from the placode contributing to fate-determination of the pre-Dc, mesenchymal expression of bone morphogenetic protein (*BMP4*) and noggin (*NOG*) to inhibit hair formation in IFE cells, and PDGFA/TGFβ signalling for hair follicle downgrowth^21^ (**Extended Data Fig. 4j**). Additionally, similar to fibroblast differentiation in murine skin, the pre-Dc, Dc, Dp and dermal fibroblasts in human prenatal skin also originated from a common fibroblast progenitor (*HOXC5^+^* early fibroblast) (**Fig. 2f**). However, dermal fibroblast differentiation into histologically defined subsets (papillary and reticular) appears to occur earlier in mice (∼E12.5)^67^. Our human prenatal skin fibroblasts did not significantly express known markers of adult human papillary fibroblasts (*COL13A1*, *COL23A1*, *PTGDS*, *ENTPD1*)^68–70^ (**Extended Data Fig. 4k)**, suggesting that the distinction between papillary and reticular fibroblasts emerges after 16 PCW. This may be attributed to differences in gestation lengths and tempo of differentiation between human and mouse. Cellular differentiation to form different tissues during development occurs at a quicker pace in mice compared to humans^71–73^ but the longer human gestation permits more advanced maturation to take place *in utero.* In keeping with this, mouse organs are less mature at birth and newborn mouse skin, which lacks hair, physically resembles early second trimester human skin.

### Genetic hair and skin disorders

Having mapped the formation and differentiation of human prenatal skin and skin organoid hair follicles, we leveraged this information to gain insights into genetic hair disorders, and to understand the extent to which hair and skin diseases have their roots *in utero*. Mutations in genes known to cause reduced hair growth (hypotrichosis) or abnormally-shaped hair (pili torti, woolly and uncombable hair syndromes) (**Supplementary Table 9**) were found to be expressed along the ‘ORS/CL’, ‘IRS’ and ‘hair fibroblast’ trajectory pseudotimes (**Extended Data Fig. 4b, 5a, b**) and in prenatal hair follicle cell states (**Extended Data Fig. 5c**), suggesting that these disorders result from dysfunctional hair follicle development.

We assessed the expression of genes causing epidermolysis bullosa (EB), an inherited blistering skin disorder that presents at birth or during early infancy, reflecting *in utero* onset of the disease^74^. In keeping with the clinical features of skin fragility secondary to structural defects in the epidermis and adjacent dermoepidermal junction, we observed higher expression of genes implicated in EB in epidermal compared to dermal prenatal skin cells (**Extended Data Fig. 5d**). Gene therapy studies for dystrophic EB have identified fibroblasts expressing *COL7A1* as a promising therapeutic strategy^75,76^. We observed *COL7A1* expression across several fibroblast subsets within the dermal compartment of prenatal tissue and skin organoids, lending support to gene therapy approaches. The expression of genes implicated in congenital ichthyoses, a group of disorders due to abnormal epidermal differentiation^77^, were primarily confined to keratinocytes (**Extended Data Fig. 5e**).

We observed similar expression patterns across prenatal skin and skin organoid for the above genetic hair and skin disorders (**Extended Data Fig. 5c-e**), supporting the value of skin organoids as a model to study congenital diseases. Although the expression of genes implicated in these disorders are confined to structural cells, disease manifestations are often associated with immune infiltration, implicating skin-immune crosstalk during pathogenesis^78,79^.

### Early dermal fibroblasts and macrophages protect skin against scarring

Prenatal human skin is uniquely able to heal without scarring but loses this capacity after 24 PCW^80,81^. Scars result from aggregation of collagen produced by dermal fibroblasts and failure of the overlying epidermis to completely regenerate^82^. To identify the cellular and molecular mechanisms that may endow early prenatal skin with scarless healing properties, we investigated the temporal changes in composition and transcriptional profile of the dermal fibroblast subsets (*FRZB*^+^, *HOXC5*^+^, *WNT2*^+^, *PEAR1*^+^ fibroblasts) (**Extended Data Fig. 3b, 6a**). We first compared prenatal skin dermal fibroblasts with healthy adult skin fibroblasts (annotated F1-F3 as in published data)^13^. All adult fibroblast subsets expressed higher levels of inflammatory cytokines (*IL-6*, *IL32*) and receptors (*IL1R1*, *IFNGR1, IL15RA, OSMR*), interferon-induced genes (*IFITM1*) and genes involved in antigen presentation (*HLA-B*, *HLA-E),* complement pathway and innate immune response (*C1R*, *C1S*, *CD55*), inflammatory response (*PTGES*, *SQSTM1*), lipid metabolism (*APOD*, *ARID5B*) and cellular senescence (*CDKN1A*) (**Fig. 3a**). In contrast, prenatal skin fibroblasts upregulated genes involved in immune suppression (*CD200*), tissue remodelling (*FAP*) and immune evasion, which are also expressed by cancer-associated fibroblasts (*THY1*, *CXCL12*, *TGFBI*)^83–86^ (**Fig. 3a**).

**Fig. 3:**
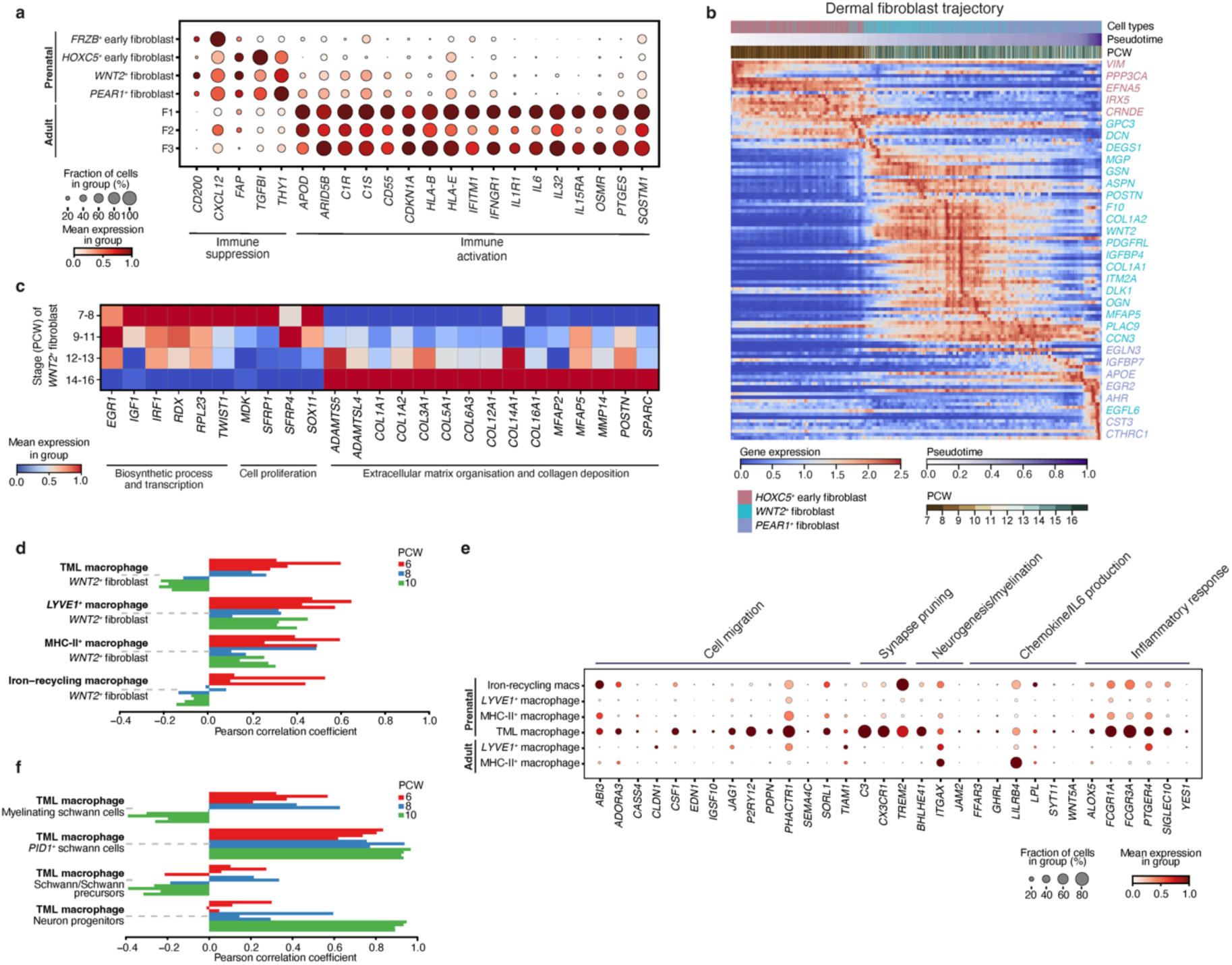
Early dermal fibroblasts and macrophages protect against skin scarring. **(a)** Dot plot showing variance-scaled, mean expression (dot colour) and percent of expressing cells (dot size) of genes differentially expressed by prenatal (‘immune suppression’) and adult skin fibroblasts (‘immune activation’)^13^. **(b)** Heat map showing differentially expressed genes across pseudotime along the ‘Dermal fibroblast trajectory’. **(c)** Matrix plot showing variance-scaled, mean expression (colour) of Milo-generated DEGs by gestational age (grouped by PCW) in *WNT2^+^* fibroblast population. DEGs are grouped by function. **(c)** Bar plot showing cell type co-location, indicated by positive Pearson correlation coefficient, for selected cell type pairs (macrophages and *WNT2^+^* fibroblasts). Pearson correlation coefficients were calculated across all skin-covered spots of Visium samples; each sample is shown by an individual bar. **(e)** Dot plot showing variance-scaled, mean expression (dot colour) and percent of expressing cells (dot size) of genes differentially upregulated by TMLM in prenatal and adult skin macrophages. Genes are grouped by function. **(f)** Bar plot showing cell type co-location, indicated by positive Pearson correlation coefficient, for selected cell type pairs (TMLM and neuronal cells). Pearson correlation coefficients were calculated across all skin spots of Visium samples; each sample is shown by an individual bar. TML macrophage: *TREM2^+^*microglia-like macrophage.

The adult fibroblast gene expression profile (‘immune activation’) was increased in *WNT2^+^* and *PEAR1^+^* dermal fibroblasts, which were abundant in later gestation, compared to *FRZB^+^*and *HOXC5^+^* fibroblasts found in earlier gestation (**Fig. 3a, Extended Data Fig. 3b, 6b**). Genes associated with a pro-inflammatory fibroblast phenotype (*APOE*, *IGFBP7, ITM2A*)^69,87^ were also upregulated during the transition from *HOXC5^+^* fibroblasts into *PEAR1*^+^ fibroblasts in prenatal skin as seen along the ‘dermal fibroblast’ pseudotime (**Fig. 3b**). In addition to transcriptomic heterogeneity between fibroblast subsets enriched in early versus late gestation, Milo^18^ analysis also revealed differences within the *WNT2^+^* fibroblast population across gestation time (**Extended Data Fig. 1d**). Late gestation *WNT2^+^* fibroblasts upregulated function-defining genes related to extracellular matrix and collagen deposition (e.g., *COL1A1*, *MFAP2*, *POSTN*), while early *WNT2^+^* fibroblasts differentially expressed regulators of transcription (*EGR1*, *IRF1*) and cellular proliferation (*SFRP1*, *SOX11*) (**Fig. 3c, Extended Data Fig. 6c, Supplementary Tables 10-12**).

Notably, expression levels of several genes involved in cellular senescence (*CDKN1A)*, cytokine pathways (*IL1R1*, *IL32),* and collagen deposition (*COL1A1*, *COL5A1*, *POSTN*), which were upregulated in *WNT2^+^* and *PEAR1^+^* prenatal fibroblasts (**Fig. 3a-c**), are also increased in pathogenic fibroblasts of fibrotic skin disorders, such as systemic sclerosis and keloid scars^88–90^. This further supports our finding of progressive acquisition of scar-promoting genes in later gestation, consistent with the clinical observation of scarring in third trimester skin^91,92^.

The role of macrophages in promoting wound healing has been described in postnatal murine and adult human skin^93–95^. In prenatal skin, we observed macrophage subsets (**Extended Data Fig. 6d, e**) co-locating with fibroblasts, neuronal and vascular cells in distinct tissue microenvironments in early gestation (**Fig. 1d, e**). Specifically, *LYVE1^+^*macrophages co-located with *WNT2*^+^ fibroblasts and were predicted to interact through platelet-derived growth factors (PDGF) with corresponding receptors (PDGFRα/PDGFRβ) expressed on fibroblasts (**Fig. 3d, Extended Data Fig. 6f, Supplementary Table 7**). Interactions between macrophages and fibroblasts maintain tissue homeostasis in diverse organs such as spleen, peritoneum and heart^96^. Our identification of additional growth factor interactions (*IGF1*-*IGF1R, GRN-EGFR*) (**Extended Data Fig. 6f, Supplementary Table 7**) suggests a role of *LYVE1*^+^ macrophages in maintenance of prenatal skin dermal fibroblasts.

We recently identified yolk-sac derived *TREM2*^+^ macrophages that share an expression profile with microglia-like macrophages from other developing organs, such as the brain, prenatal skin and gonads^97–99^. *TREM2*^+^ microglia-like macrophages (TMLM), including those in prenatal skin, expressed *P2RY12*, *CX3CR1*, *TMEM119*, *OLFML3*, *C3* (**Extended Data Fig. 6d, e**). Prenatal skin TMLM were highly correlated with embryonic brain microglia^97^ (**Extended Data Fig. 7a, b**) and co-expressed immunomodulatory genes, including immune-inhibitory receptors (*LILRB4*, *SIGLEC10*, *CX3CR1*) and regulators of IL-6 production (*SYT11*, *GHRL*)^100–102^ (**Fig. 3e, Extended Data Fig. 7c, Supplementary Tables 13-17**). Downregulation of inflammation and IL-6, secreted paracrinally by macrophages and autocrinally by fibroblasts^96^, has been reported to confer anti-fibrogenic properties in mouse skin transplants and fetal wounds^103,104^. Furthermore, blocking of IL-6 receptor resulted in decreased production of pro-fibrotic molecules, such as collagen alpha 1, by explant dermal fibroblasts from systemic sclerosis patients^105^. TMLM co-located with *WNT2*^+^ fibroblasts in early prenatal skin (6-8 PCW) (**Fig. 3d)** and *WNT2*^+^ fibroblasts downregulated *IL6* expression compared to adult fibroblasts (**Fig. 3a**). This suggests a potential contribution to scarless healing in prenatal skin by TMLM through interactions with *WNT2*^+^ fibroblasts. Accordingly, *GAS6*, expressed by TMLM, was predicted to interact with *TYRO3*/*AXL* receptors on *WNT2*^+^ fibroblasts (**Extended Data Fig. 6f, Supplementary Table 7**) and these interactions have been shown to induce immunosuppression, resolution of inflammation, and tissue repair in the synovium, lung, and liver^106–108^.

Collectively, our findings suggest that the composition and gene expression profile of prenatal skin fibroblasts in early gestation favour tissue remodelling over extracellular matrix formation and collagen deposition. Our data also suggests a role for TMLM, which express immunoregulatory gene profile and co-locate with *WNT2*^+^ fibroblasts, in conferring the unique property of scarless healing observed in early prenatal skin.

### Macrophages in cutaneous neuronal differentiation

TMLM in prenatal skin are also co-located with neuronal Schwann cells (‘neurovascular microenvironment’) (**Fig. 1d**, **3f**) and expressed genes related to cell migration and neuronal development (**Fig. 3e, Extended Data Fig. 7c, Supplementary Tables 13, 14**), mirroring the functions of brain microglia and peripheral nerve-associated macrophages in mouse skin^109–112^. TMLM were predicted to interact with Schwann cells (CellPhoneDB), contributing to synapse formation and axon guidance (*VEGFA*-*NRP1/2*, *SEMA3C*-*NRP2, SEMA3E*-*PLXND1*) (**Extended Data Fig. 7d, Supplementary Table 7**)^113–115^. These findings suggest that dermal macrophages may support neuronal development in prenatal skin, including a specific contribution of TMLM to the establishment of the skin peripheral nervous system during early gestation as previously reported in murine skin^110,111,116^.

### Macrophages support prenatal skin angiogenesis

Macrophages have been implicated in angiogenesis during prenatal organ development including the heart, aorta-gonad-mesonephros and brain, as well as in the post-natal setting such as cancer-related angiogenesis^9,117–119^. Blood vessel formation is critical for nutrient and oxygen support during tissue morphogenesis. Tissue primary vascular beds arise following the differentiation of primitive mesoderm cells into endothelial progenitors which migrate and coalesce (vasculogenesis)^120^. Endothelial cells can proliferate or acquire a specialised ‘tip’ phenotype to guide blood vessel branching and network formation (sprouting angiogenesis) in response to pro-angiogenic signals^121^.

We observed close proximity of all four prenatal skin macrophage subsets (iron-recycling, *LYVE1^+^*, MHC-II*^+^* and TMLM) to endothelial cells by Visium deconvolution analysis (‘neurovascular microenvironment’) (**Fig. 1d, e)**, which was further validated by multiplex RNAScope staining (**Fig. 4a**). The four macrophage subsets expressed gene programmes driving angiogenesis (**Extended Data Fig. 7c, Supplementary Tables 13-17**). We sought to investigate if these macrophage subsets contribute to distinct angiogenic processes using module scoring of reference angiogenesis gene sets^122,123^ and cell-cell interaction analysis (**Extended Data Fig. 7e, 8a, Supplementary Tables 18, 19**). Gene module expression profiles suggested that sprouting angiogenesis (growth of new vessels) was promoted by *LYVE1^+^*and MHC-II*^+^* macrophages, endothelial cell chemotaxis by iron-recycling macrophages, and endothelial cell proliferation and blood vessel morphogenesis by *LYVE1^+^* macrophages (**Extended Data Fig. 7e, Supplementary Table 18**). Predicted ligand-receptor interactions showed the reciprocal communication between macrophages and endothelial cells supporting angiogenesis, chemotaxis, and cell migration (**Extended Data Fig. 8a, Supplementary Table 19**). We identified *CXCL8* and *CCL8* on *LYVE1^+^* macrophages, interacting with *ACKR1* on vascular endothelial cells to regulate angiogenesis, as previously reported (**Extended Data Fig. 8a, Supplementary Table 19**)^13,124^.

**Fig. 4:**
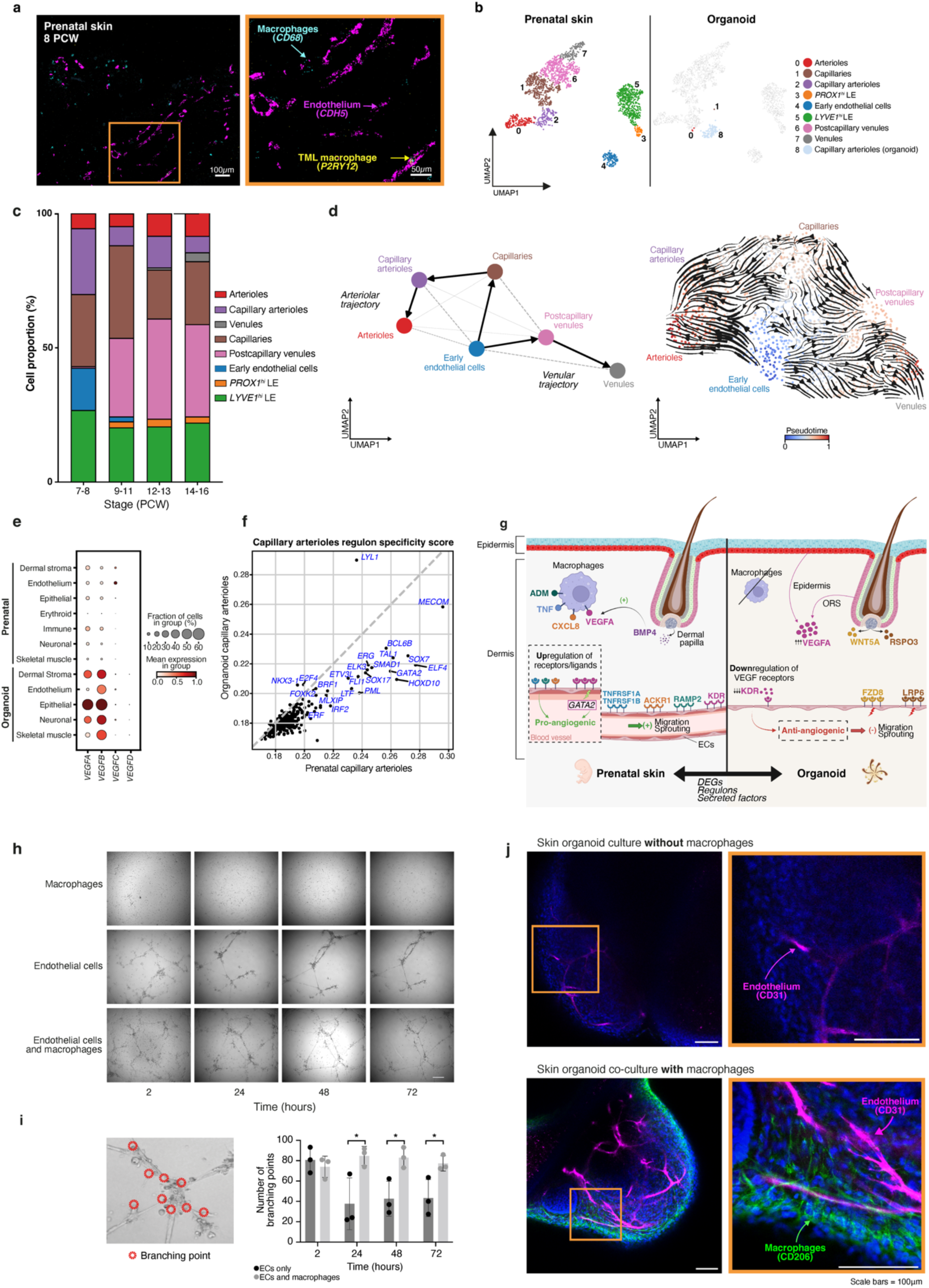
Macrophages support prenatal skin angiogenesis. **(a)** Large-area (left, scale bar = 100μm) and magnified (right, scale bar = 50μm) RNAScope images of prenatal skin (representative 8 PCW sample) demonstrating endothelium (*CDH5*), macrophages (*CD68*) and TMLM (*P2RY12*). **(b)** UMAP visualisation of endothelial cell states present in prenatal skin and skin organoid. **(c)** Bar plot showing the proportions of endothelial cell states across gestational age in prenatal skin. Bar colours represent cell states. **(d)** Inferred pseudotime trajectory of prenatal skin endothelial cell states differentiating along ‘arteriolar’ and ‘venular’ trajectories; UMAP overlaid with PAGA and coloured by cell state (left) and overlaid with CellRank state transition matrix inferred arrows and coloured by pseudotime (right). **(e)** Dot plot showing variance-scaled, mean expression (dot colour) and percent of expressing cells (dot size) of vascular endothelial growth factors in prenatal skin and skin organoids. **(f)** Comparison of regulon activity between prenatal skin (x-axis) and organoid (y-axis) capillary arterioles. **(g)** Schematic showing differences between prenatal skin and skin organoids in pro- and anti-angiogenic factors produced by myeloid and dermal papilla cells and their corresponding receptors on endothelial cells. **(h)** Angiogenesis assay including macrophages only, endothelial cells only and co-culture of endothelial cells plus macrophages at timepoints 2 hours, 24 hours, 48 hours, and 72 hours of culture. Scale bar=200μm. **(i)** Analysis of the number of branch points in each condition of the angiogenesis assay. **(j)** Representative wholemount immunofluorescence images of the skin organoid, without and with macrophage co-culture, showing macrophages (CD206) in green, endothelium (CD31) in red and DAPI nuclei stain in blue. Scale bars=100μm. DEGs: differentially expressed genes, EC: endothelial cells, LE: lymphatic endothelium, TML macrophage: *TREM2^+^* microglia-like macrophage.

Our data suggested macrophages contribute to prenatal skin angiogenesis. Consistent with this observation, in skin organoids which lacked immune cells, vascularisation was poor despite formation of well-developed hair follicles, epidermis and neuronal cells (**Fig. 4b**). Furthermore, almost all endothelial cells in the skin organoid only expressed markers of capillary arterioles over the developmental time course^125–127^ (**Fig. 4b, Extended Data Fig. 8b, Supplementary Table 20**). In contrast, prenatal skin was populated by heterogeneous endothelial cells, including early endothelial cells before 9 PCW, followed by a predominance of capillaries, post-capillary venule and lymphatic endothelial cells after 9 PCW (**Fig. 4 b-c**).

Inferred trajectory analysis of prenatal skin vascular endothelium showed that early endothelial cells differentiated into either an arteriolar (capillaries, capillary arterioles, arterioles) or venular (postcapillary venules and venules) pathway (**Fig. 4d, Extended Data Fig. 9a**). This was supported by the differential expression across pseudotime of known genes characterising arteriolar (*AQP1*, *GJA5*, *FBLN5*, *CXCL12*) and venular (*ACKR1*, *PLVAP*, *CLU*) endothelial cells^125–127^ (**Extended Data Fig. 9b, c**). Our trajectory analysis suggested that prenatal skin capillary arteriole cells possessed the capacity to differentiate further into arterioles (**Fig. 4d, Extended Data Fig. 9a**), unlike organoid capillary arteriole cells. Additional comparison of prenatal skin with a human ES/iPS-derived blood vessel organoid^128^, which also lacked immune cells, predicted endothelial cells from this model to be early endothelial, capillary arteriole, or arteriole cells by logistic regression analysis (**Extended Data Fig. 9d**). This demonstrates limited vasculature differentiation even in a mesoderm-geared blood vessel organoid model and is consistent with a requirement for immune cells to fully recapitulate the endothelial cell development observed in prenatal skin.

We investigated additional potential mechanisms for failed expansion and differentiation of skin organoid endothelial cells. We considered whether recognised drivers of angiogenesis^129^ were absent, including laminar blood flow, hypoxia, and the sprouting angiogenesis potential of capillary arteriole cells. Indeed, genes activated by blood flow were upregulated in capillary arteriole cells in prenatal skin but not in the skin organoid (**Extended Data Fig. 9e**). However, the gene module scores of hypoxia-related genes (**Supplementary Table 20**) were reduced in the skin organoid relative to prenatal skin for corresponding cell categories (**Extended Data Fig. 9f**). The potential for sprouting angiogenesis (characterised by the ‘tip’ cell state) was assessed using the ‘tip’ score for each of the endothelial cell subsets in prenatal skin and skin organoid^130^. Prenatal skin arteriole, capillary arteriole, and capillary cells, as well as organoid capillary arteriole cells had increased ‘tip’ scores, reflecting a high sprouting angiogenesis capability (**Extended Data Fig. 9g, h**). This suggests that despite strong expression of the sprouting angiogenesis gene signature, organoid capillary arteriole cells were unable to guide stalk cells for new blood vessel formation.

We next compared the expression profile of angiogenesis-related genes in prenatal skin and skin organoid. Several pro-angiogenic genes (*CXCL8*, *TNF*, *ADM*), and corresponding receptors, were upregulated in prenatal skin compared to the skin organoid and were primarily expressed by macrophages (**Extended Data Fig. 10a, Supplementary Tables 21-25**). Conversely, skin organoid cells, such as the Dp, expressed high levels of anti-angiogenic genes, including *WNT5A* and *RSPO3*, with matching receptors on capillary arteriole cells (**Extended Data Fig. 10a, Supplementary Tables 21-25**). Vascular endothelial growth factors (VEGF), *VEGFA* and *VEGFB,* were however highly expressed in several skin organoid cells (**Fig. 4e**), but their receptors (*KDR*, *FLT1*) on capillary arterioles were downregulated in the organoid compared to prenatal skin (**Extended Data Fig. 10b**). Regulon analysis (pySCENIC^131^) showed downregulated *GATA2* and its target regulons (*MECOM*, *BCL6B*, *SOX7* and *ELF4*) in skin organoid capillary arterioles compared to prenatal skin (**Fig. 4f, Extended Data Fig. 10c**). *GATA2* plays a key role in angiogenesis during development and has been shown to regulate VEGF-induced endothelial cell migration and sprouting *in vitro*^132–134^. Several target genes of *GATA2* and *SOX7* (*APLNR*, *AQP1*, *CAV1*, *VWF*) (**Extended Data Fig. 10d**) were expressed across the ‘arteriolar’ and ‘venular’ trajectory pseudotimes in prenatal skin (**Extended Data Fig. 9b, c**) and are involved in endothelial cell differentiation^125^. These genes were downregulated in the skin organoid capillary arterioles compared to prenatal skin (**Extended Data Fig. 10b**). VEGF receptors (*KDR* and *FLT1*), which were downregulated in the skin organoid, were also inferred to be downstream targets of *GATA2* as previously reported^133,135^ (**Extended Data Fig. 10d**). An orthogonal approach using NicheNet^136^ identified similar macrophage-expressed pro-angiogenic factors, such as *TNF*, as driving differential gene expression between prenatal skin and skin organoid endothelial cells (**Supplementary Tables 26-28**). This also identified *VEGFA* as one of the top upstream ligands regulating differences in *GATA2* expression (**Extended Data Fig. 10e, Supplementary Tables 26-28**). These findings suggest that high VEGF signalling in the skin organoid cannot compensate for missing macrophage-related factors in driving *GATA2* activity and downstream VEGF receptor expression (**Fig. 4g**).

The above data implicates prenatal skin macrophages as key players in angiogenesis, supporting our earlier observation of pro-angiogenic gene module expression in TMLM in the human yolk sac^99^. We next set out to validate our findings using two distinct cell culture assays. We performed an angiogenesis assay by co-culturing iPS-derived endothelial cells with and without iPS-derived macrophages (both differentiated from the iPS cell line Kolf2.1S) (**Extended Data Fig. 10f**). The macrophage differentiation method used here has been shown to recapitulate features of primitive and definitive hematopoiesis^137^. Endothelial cells, generated according to a previously published protocol^138^, were cultured alone or in co-culture with macrophages and began network formation with similar numbers of branching points measured at 4 hours of culture (**Fig. 4h, i**). The number of branches dropped significantly in the endothelial-only culture by 24 hours but was maintained in the presence of macrophages up to 72 hours (**Fig. 4h, i**).

We then tested whether the limited angiogenesis observed in skin organoids could be rescued by supplying missing immune cells. We introduced iPS-derived macrophages in the early stages of the skin organoid differentiation and assessed the endothelial network in skin organoid by whole mount immunofluorescence imaging on day 35 of co-culture, using CD206 to identify mature macrophages and CD31 for endothelial cells. Macrophages co-localised with blood vessels, even after 5 weeks of culture (**Fig. 4j, Extended Data Fig. 10g**). We observed a more elaborate and developed vascular network in skin organoids co-cultured with macrophages compared to control organoids without macrophages (**Fig. 4j, Extended Data Fig. 10g**). These *in vitro* findings in a 2D angiogenesis assay and 3D skin organoid culture model demonstrate that cell-cell interactions between macrophages and endothelial cells provide the necessary cues to support angiogenesis and improve blood vessel expansion.

## Discussion

In this study, we characterise the dynamic composition of human prenatal skin during the early stages of *de novo* hair follicle formation and highlight the critical skin immune and non-immune crosstalk contributing to skin morphogenesis. We provide evidence for the contributions of immune cells to human prenatal skin development, in line with emerging evidence from tissue morphogenesis in animal studies^139^. Notably, macrophage subsets appear to play distinct roles in scarless skin repair, fibroblast homeostasis and neurovascular development. This is in part contributed to by yolk-sac derived TMLM^99,140^, which have also been reported in human prenatal testicular cords, suggesting there is a broader tissue distribution in early gestation than previously thought and functions outside of the central nervous system.

We identified a critical role of macrophages in angiogenesis which could not be compensated for by culture conditions or exogenous growth factors *in vitro*. Adding macrophages to endothelial cell cultures and hair-bearing iPS-derived skin organoids enhanced endothelial cell numbers and vascular network formation. Notably, iPS-derived macrophages survived at least five weeks in the skin organoid, in contrast to 1-2 weeks in liquid culture supplemented with macrophage colony-stimulating factor (M-CSF)^137^, highlighting the importance of tissue microenvironment in sustaining macrophage residency and longevity. The development of other structures such as hair follicles, melanocytes, neuronal and interfollicular epidermal and dermal cells in the skin organoid were unaffected by the absence of immune cells. This is likely due to the specific timing and supplementation of growth factors to promote skin and hair follicle differentiation in the hair-bearing skin organoid model. Indeed, the requirement for macrophages for adequate development of both ectoderm and mesoderm-derived structures, including peripheral neuronal cells and blood vessels, has been demonstrated in mouse models of macrophage depletion^141–144^.

Our study provides further insights into human hair follicle formation and the origin of the companion layer, which appears to develop along the same trajectory as the outer root sheath, and not from the hair matrix, as previously reported^21,145,146^. A recent murine fate-mapping study also showed that the CL develops prior to hair matrix formation and in the absence of matrix cells^146^. In addition, a recent single cell analysis of mouse hair follicles identified greater transcriptional similarity of CL cells to ORS cells^38^, consistent with our findings in human prenatal skin.

A combination of fibroblast and macrophage associated molecular features, including the presence of fibroblast progenitors, a down-regulated immune milieu and reduced collagen expression, are likely to contribute to the ability of prenatal skin to heal without scarring. However, although skin fibroblast differentiation is incomplete by 16 PCW, progressive ‘ageing’ is already evident with gradual acquisition of the pro-inflammatory phenotype, characteristic of adult fibroblasts^69,87^, beginning *in utero* from as early as 9 PCW. Future studies on how the microbiota colonisation and environmental exposure after birth affects skin across the human lifespan are required.

A hugely valuable utility of our prenatal human skin single cell atlas is the ability to identify expression of genes known to cause congenital hair and skin disorders, which we provide as a resource with the accompanying web portal for our study (https://developmental.cellatlas.io/fetal-skin; password: fs2023). We find that implicated genes are indeed expressed in prenatal skin, supporting an *in utero* origin for these disorders. Notably, structural cells express disease causing genes suggesting that the accompanying immune infiltration observed in these diseases results from defects in structural cells. Our study on human prenatal skin development is a valuable blueprint to guide more faithful *in vitro* skin organoid generation, which can enable future studies on interactions with the microbiota, the pathogenesis of congenital skin disorders, and hair and skin engineering for therapeutic applications, including hair regeneration and skin transplant.

## Supporting information

Combined Supplementary Tables

## Methods

### Tissue acquisition and processing

Human developmental tissue samples used for this study were obtained from the MRC– Wellcome Trust-funded Human Developmental Biology Resource (HDBR; http://www.hdbr.org) with written consent and approval from the Newcastle and North Tyneside NHS Health Authority Joint Ethics Committee (08/H0906/21+5) and Cambridge University Hospitals NHS Foundation Trust (NHS REC 96/085).

Tissues were processed into single cell suspensions immediately upon receipt for single cell transcriptomic profiling. Tissue was first transferred to a sterile 10 mm^2^ tissue culture dish and cut in <1 mm^3^ segments using scalpel. It was then digested with type IV collagenase (final concentration of 1.6 mg/ml; Worthington) in RPMI (Sigma-Aldrich) supplemented with 10% heat-inactivated fetal bovine serum (FBS; Gibco), at 37°C for 30 min with intermittent agitation. Digested tissue was then passed through a 100 μm cell strainer and cells were collected by centrifugation (500 g for 5 min at 4°C). Cells were treated with 1× RBC lysis buffer (eBioscience) for 5 min at room temperature and washed once with Flow Buffer (PBS containing 5% (v/v) FBS and 2 mM EDTA) prior to cell counting and antibody staining. Single cell suspensions were generated from skin of 15 donors with ages spanning from 7 PCW to 16 PCW.

### Single-cell RNA sequencing experiment

Dissociated cells were stained with anti-CD45 antibody (3 µL CD45 BUV395, clone: HI30, BD Biosciences) on ice in the dark for 30 min. For F69 and F71, additional staining with anti-CD34 (4 µL CD34 APC/Cy7, clone: 581, Biolegend) and anti-CD14 antibodies (3 µL CD14 PE-CF594, clone: MφP9, BD Biosciences) was also performed to improve capture of less abundant cell populations from the CD45^-^ fraction, such as keratinocytes and endothelial cells, by separating them from the abundant CD34^+^CD14^-^ stromal cells. Immediately prior to sorting, cells were passed through a 35 µm filter (Falcon) and DAPI (Sigma-Aldrich) was added at a final concentration of 3 μM. Sorting by flow cytometry was performed with BD FACSAria Fusion Flow Cytometer. CD45 positive fraction was sorted from DAPI^−^CD45^+^ gate, and CD45 negative fraction was sorted from DAPI^−^CD45^−^ gate. CD45 gating was contiguous so that no live cells were lost in sorting. Live CD45^+^ and CD45^-^ cells were sorted into separate chilled FACS tubes coated with FBS. For F69 and F71, in addition to the live CD45^+^ and CD45^-^ cells, we isolated all cells from the CD45^-^ fraction that were not within the CD34*^+^*CD14^-^ gate and collected them into a separate chilled FACS tubes coated with FBS (**Extended Data Fig. 1a).**

FACS sorted cell suspensions were counted and loaded onto the 10x Genomics Chromium Controller to achieve a maximum yield of 10,000 cells per reaction. Either Chromium single cell 3’ reagent kit (v2) or Chromium single cell V(D)J kits from 10x Genomics were used. DAPI–CD45+ or DAPI–CD45– FACS-isolated cells were loaded onto each channel of the Chromium chip following the manufacturer’s instructions before droplet encapsulation on the Chromium controller. Gene expression (GEX) and T cell receptor (TCR) libraries were generated according to the manufacturer’s instructions. The GEX libraries were sequenced to achieve a minimum target depth of 20,000 reads per cell and the TCR libraries were sequenced to achieve a minimum target depth of 5,000 reads per cell using Illumina sequencing instruments.

### Endothelial cell and skin organoid co-culture with macrophages

Endothelial cells were derived from Kolf2.1S iPS cells (iPSCs) cultured on matrigel coated plates in mTeSR1 medium with ROCK inhibitor at 4.5x10^4^ cells per cm^2^. iPSCs were differentiated through lateral mesoderm into CD144 positive endothelial cells as previously published^138^. Macrophages and skin organoids were also derived from Kolf2.1S iPSCs according to previously published methods^15,137^. The angiogenesis assay was carried out by culturing iPS-derived endothelial cells and macrophages separately or in co-culture in 15-well 3D chambered µ-slide (ibidi GmbH, cat. no. 81506). This was done using a three-layered sandwich method where layer one was 10µl matrigel (Corning, cat. no. 354230), layer two was supplemented StemPro medium (Gibco, cat no. 10639011) + 10% matrigel with and without the endothelial cells and layer three was supplemented StemPro medium with and without macrophages. The endothelial cells were left to settle for 4 hours at 37°C before addition of macrophages. The assay was imaged 2 hours after initial culture and then 24-hourly for three days using the Olympus IX83 inverted microscope and images were analysed using ImageJ2 (v2.14.0)^147^. Prior to co-culture, iPS-derived macrophages were phenotyped using flow cytometry (**Extended Data Fig. 10f**). Macrophages were harvested using TrypLE (Gibco) at 37°C, 5% CO2 for 5 min and cells were collected by centrifugation (500 g for 6 min). Cells were washed once with Cell Staining Buffer (Biolegend) prior to cell counting and antibody staining. Dead cells were stained with Live/Dead fixable blue (ThermoFisher Scientific) for 30 min on ice in the dark. Non-specific bindings were blocked using Human TruStain FcX (Fc Receptor Blocking Solution, BioLegend) for 10 min on ice in the dark. Cells were washed twice with Cell Staining Buffer. Single-staining was performed on cells with anti-CD206 antibody (1:20 CD206 PE, clone 19.2, ThermoFisher Scientific), anti-CD16 antibody (1:20 CD16 PE-Cyanine7, clone eBioCB16, ThermoFisher Scientific), anti-CD14 antibody (1:20 CD14 PerCP-Cyanine5.5, clone 61D3, ThermoFisher Scientific) and anti-CD1c antibody (1:25 CD1c Pacific Blue, clone L161, BioLegend) on ice in the dark for 30 min. Prior to acquiring on analyzer, cells were washed once in Cell Strained Buffer and passed through a 35 µm filter (Falcon). Acquisition by flow cytometry was performed with BD LSRFortessa Cell Analyzer. Live single CD16^+^, CD14^+^, CD206^+^ and CD1c^+^ cells were analysed using FlowJo (version 10.9.0). The co-culture was performed by adding the macrophages to the skin organoids on day 12 of culture, with a 1:5 ratio. Skin organoids were transferred to a low attachment 96-well plate (Nunclon Sphera, Life Technologies) in skin organoid maturation medium^15^ containing 10% or 20% matrigel (Corning). 2000 macrophages were added to the skin organoids and the co-culture was centrifuged at 100g for 3 min 1acc, 0dec. Medium was changed every other day. On day 8 of co-culture, the cells were transferred to a low attachment 24-well plate and matrigel was diluted with fresh skin organoid maturation medium. On day 35 of co-culture, the cells were fixed in 2ml tube with 4% PFA overnight. The co-culture was then permeabilized in blocking buffer (0.3% (vol/vol) Triton X-100, 1% (vol/vol) normal goat serum based on the antibodies and 1% BSA (vol/vol) dissolved in 1× PBS) for 4 hours at RT. Cells were then incubated overnight at 4°C on a shaker (65 rpm) with primary antibodies as follows: anti-CD206 (Abcam) for macrophages and anti-CD31 (Dako) for endothelial cells. The morning after, cells were washed and then incubated with secondary antibodies and DAPI for 4 hours on a shaker at RT as follows: goat anti-rabbit IgG, Alexa Fluor Plus 488 and goat anti-mouse IgG1, Alexa Fluor 568 (ThermoFisher). Cells were washed and placed in 50% glycerol for 30 minutes on a shaker at RT. Cells were then transferred to 70% glycerol overnight on a shaker at RT. The following morning, the co-culture was mounted and imaged using the Leica SP8 microscope.

### Visium spatial data generation

Prenatal facial (n=1, replicate=2) and abdominal skin (n=1, replicate=2) samples from a single donor at 10 post conception weeks (PCW) were embedded in optimal cutting temperature (OCT) medium and flash-frozen in isopentane cooled with dry-ice. 10µm cryosections from the OCT blocks were cut onto 10x Genomics Visium slides. Sections were stained with hematoxylin and eosin and imaged at 20X magnification on a Hamamatsu Nanozoomer. These sections were then processed according to the 10x Genomics Visium protocol, using a permeabilization time of 12 min found through a prior tissue optimization step. Dual-indexed libraries were prepared as per manufacturer’s protocol, pooled at 2.8 nM and sequenced in 8 samples per Illumina Novaseq S4 flow cell with the following run parameters: read 1: 28 cycles; i7 index: 10 cycles; i5 index: 10 cycles; read 2: 90 cycles.

### Multiplex RNAScope staining and imaging

Prenatal skin tissue (8 and 15 PCW) was frozen in OCT compound (Tissue-Tek O.C.T). 4-plex smFISH was performed using the RNAScope Multiplex Fluorescent Detection Kit v2 (ACDBio, Newark, California, USA, cat. no. 323100) according to the manufacturer’s instructions using the standard pre-treatment for fresh frozen sections of 10-20 μm and permeabilized with Protease IV, for 30 mins at room temperature.

Human probes against FOXP3, SHH, SLC26A7, NDP, CDH5, CD68, P2RY12 mRNA molecules were used (all from ACDBio catalogue probes, Newark, California, USA). Opal dyes (Akoya Biosciences, Marlborough, Massachusetts, USA) were used at a dilution of 1:1,000 for the fluorophore step to develop each channel: Opal 520 Reagent Pack (FP1487001KT), Opal 570 Reagent Pack (FP1488001KT) and Opal 650 Reagent Pack (FP1496001KT) and Atto-425. Finally, the slides were counterstained with DAPI and coverslipped for imaging with ProLong Gold Antifade Mountant (ThermoFisher, Canoga Park, California, cat. no. P36930).

Confocal imaging was performed on a Perkin Elmer Opera Phenix Plus High-Content Screening System using a 40X (NA 1.1, 0.149 μm/pixel) water-immersion objective with a 2µm z-step. Channels: DAPI (excitation 375 nm, emission 435-480 nm), Atto 425 (ex. 425 nm, em. 463-501 nm), Opal 520 (ex. 488 nm, em. 500-550 nm), Opal 570 (ex. 561 nm, em. 570-630 nm), Opal 650 (ex. 640 nm, em. 650-760 nm). Confocal image stacks were stitched as two-dimensional maximum intensity projections using proprietary Acapella scripts provided by Perkin Elmer and visualised using OMERO Plus (Glencoe Software).

### scRNA-seq data analysis

#### Alignment, quality control, clustering and annotation of prenatal skin dataset

The gene expression data were mapped with CellRanger 2.1.1 and 2.0.2 to an Ensembl 84– based GRCh38 reference (10X Genomics–distributed 1.2.0 version). The python package emptydrops (v0.0.5) was used to detect cells in each sample. Potential doublets were flagged by Scrublet (v0.2.1)^148^ as previously implemented^149^. Low-quality cells were filtered out [minimum number of genes = 100, maximum number of UMIs = 45000, maximum mitochondrial UMI fraction = 0.15]. Possible maternal contamination was identified using the souporcell pipeline (v2.4.0)^150^ as previously described^9^. Data pre-processing was performed using scanpy (v1.4.3)^151^. After pooling data from all samples, genes detected in fewer than 3 cells were removed, and data was normalised to 1e4 UMI per cell and log1p transformed.

Highly variable genes were selected based on normalised dispersion (scanpy.pp.highly_variable_genes with flavor = “seurat”, min_mean = 0.0125, max_mean = 3, min_dispersion = 0.5). Dimensionality reduction was done by Principal Component Analysis (PCA) and the first 50 principal components (PCs) were used to compute the nearest-neighbour graph (scanpy.pp.neighbors with n_neighbors = 15). BBKNN (v1.3.3)^152^ was used to generate a batch-corrected nearest-neighbour graph considering each donor as a separate batch. Leiden algorithm was used to cluster cells based on the corrected graph with a relatively low resolution (scanpy.tl.leiden with resolution = 0.3) into coarse clusters which were manually annotated into broad lineages using known marker genes.

For each broad lineage, the data was re-processed starting from highly variable gene selection to better reveal the finer heterogeneity. At this level, we used Harmony (v0.0.5)^153^ and BBKNN (v1.3.3)^152^ in parallel for batch correction (again treating each donor as a separate batch and with batches under 10 cells removed) for every broad lineage and observed highly consistent embedding and clustering (data provided on portal). For fibroblasts, we continued our analysis with embedding and clustering downstream of BBKNN and for all other broad lineages, we continued with Harmony. Leiden clusters at the highest resolution were annotated manually using marker genes identified through literature search and their alignment with differentially expressed genes (DEGs) in each cluster (**Supplementary Table 2**). DEGs were calculated using the sctk (Single Cell analysis Tool Kit) package (https://github.com/Teichlab/sctk), where filtering is carried out followed by a two-sided Wilcoxon rank-sum test using pass-filter genes only in a one-vs-all fashion. The sctk package also carries out comparisons between the group of interest (one with highest expression) and the next group (second highly expressed), where the maximum proportion of cells expressing the gene in question in the second most highly expressed group was 0.2. For epidermal annotations, we created a combined embedding of prenatal skin and skin organoid data^15^, integrated using the Harmony pipeline, as described below. Harmony corrected PCs were used to compute the batch-corrected nearest neighbourhood graph, and the Leiden algorithm was used to cluster the integrated data. The sctk package was then used to derive marker genes for derived Leiden clusters. Annotation was carried out on the clusters based on marker genes and refined annotations in the skin organoid data^15^.

Clusters of doublets were manually flagged and removed taking into account markers genes and previously calculated scrublet scores. To have a final global visualisation of the atlas, a doublet-free UMAP was generated (**Fig. 1b**).

#### Processing, clustering and annotation of skin organoid dataset

Organoid data was pre-processed, filtered, clustered, and annotated separately before integration with prenatal skin. Briefly, cells filtered by CellRanger (CellRanger 2.1.0 with GRCh38-1.2.0 and CellRanger 3.0.2 with GRCh38-3.0.0) from skin organoid samples (2 strains, each with four time points) were pooled and quality control (QC) thresholds for UMI counts, gene counts, percentage of mitochondrial (MT) genes and top 50 highly expressed genes were established by fitting Gaussian mixture models to the distribution of each metric respectively. The thresholds used were: minimum number of genes = 450, maximum number of genes = 5731, minimum number of UMIs = 1063, maximum number of UMIs = 25559, maximum mitochondrial UMI fraction = 0.133, minimum cumulative percentage of counts for 50 most expressed genes in a cell = 23.7%, maximum cumulative percentage of counts for 50 most expressed genes in a cell = 56.6%. Highly variable genes selection, dimensionality reduction and KNN graph construction were done using the same method and parameters as prenatal skin. BBKNN (v1.3.3)^152^ was again used for batch-correction treating combinations of strains and 10X kit versions as batches. Broad lineages were annotated based on known markers. Each broad lineage was then re-processed in the same way as prenatal skin to annotate cell types at higher resolution (**Supplementary Table 2**).

#### Integration of prenatal skin and skin organoid datasets

To integrate prenatal skin cells with organoid cells, the datasets were downsampled for each broad lineage to have roughly balanced cell counts per cell type before integration with Harmony (v0.0.5)^153^, treating datasets as batches (prenatal skin or organoid) and within dataset batches as covariates (donor for prenatal skin and strain:10X kit version for organoid). Leiden clusters were annotated using known markers.

#### Comparison of prenatal skin, adult skin and skin organoid datasets: distance-based analysis

To integrate prenatal skin, adult skin and skin organoid cells, all datasets were downsampled (max 500 cells per broad cell type) to have roughly balanced cell counts per broad lineage before integration with Harmony (v0.0.5)^153^, treating datasets as batches and within-dataset batches as covariates (donor for prenatal and adult skin and strain for organoid). The principal component vectors of the downsampled Harmony-integrated object were then used to transform the gene expression matrix (NumPy (v1.23.4) function ‘linalg.lstsq’, rcond = ‘warn’) of all cells in the non-downsampled pooled data and project for UMAP-visualisation (**Fig. 1f, Extended Data Fig. 2a**). The median transformed gene expression was used to compute the Euclidean distance between prenatal skin, adult skin and skin organoid for each broad cell cluster, using ‘spatial.distance_matrix’ function in SciPy (v1.9.3), which was then plotted as heatmap (**Extended Data Fig. 2c**).

#### Time-encoded cell state predictions: prenatal skin, adult skin and skin organoid datasets

Median probability of class correspondence between gene expression matrices in single cell datasets were carried out using a logistic regression (LR) framework previously described^154^, based on a similar workflow to CellTypist tool^155^. Annotated raw scRNA-seq datasets (prenatal skin, adult skin and skin organoid) were first concatenated, normalised, and log-transformed. Linear variational autoencoder (VAE) latent representations were computed using the LDVAE module within scvi-tools (hidden layers=256, dropout-rate=0.2, reconstruction-loss=negative binomial) with dataset and chemistry information taken as technical covariates. ElasticNet LR models were built utilising the linear_model.LogisticRegression module in the sklearn package (v0.22). The models were trained on SCVI batch-corrected low-dimensional LDVAE representation of the training data (prenatal and adult skin) using time-encoded labels (age_cell category). Regularisation parameters (L1-ratio and alpha) were tuned using the GridSearchCV function in sklearn (v1.1.3). The test grid was designed with five l1_ratio intervals (0.05, 0.2, 0.4, 0.6, 0.8), five alpha (inverse of regularisation strength) intervals (0.05,0.2, 0.4, 0.6, 0.8) at five train-test splits and three repeats for cross-validation. The unweighted mean over the weighted mean squared errors (MSEs) of each test fold (the cross-validated MSE) was used to determine the optimal model. The resultant model was used to predict the probability of correspondence between trained time-encoded labels and pre-annotated time_encoded clusters (week of culture_cell category) in the target dataset (skin organoid). The median probability of training label assignments per predesignated class overall (all cell groups) and for individual broad cell categories were computed (**Supplementary Table 4**). For visualisation, the adult skin dataset was randomly downsampled to 10% (overall or by cell lineage) and resultant LR probabilistic relationship between labels of the training and target datasets were plotted as heatmaps (**Extended Data Fig. 2d**).

#### Differential abundance analysis

Differences in cell abundance associated with gestational age were tested using Milo (v1.0.0)^18^, correcting for CD45+ and CD45- FACS isolation strategies. We first re-embedded cells into a batch-corrected latent space with a dimension of 30 using scVI model as implemented in scvi-tools considering donor and chemistry as batches. The model was trained using the 10000 most highly variable genes which were selected based on dispersion (min_mean=0.001, max_mean=10) as previously described^99^. Where FACs correction was applied, we calculated a FACS isolation correction factor for each sample *s* sorted with gate *i* as (*fs =* log(*pi*S/Si)) where *pi* is the true proportion of cells from gate *i* and S represents the total number of cells from both gates^99^. A KNN graph of cells was constructed based on distances in the latent space and cells assigned to neighbourhoods using the milopy.core.make_nhoods function (prop=0.1). The number of cells of each cell type was then counted in each neighbourhood (milopy.core.count_nhoods). Labels were assigned to each neighbourhood based on majority voting (milopy.utils.annotate_nhoods) of cell labels by frequency (>50%). To test for differential abundance across gestational age, prenatal skin data was split into 4 age bins (7-8 PCW, 9-10 PCW, 11-13 PCW, 15-16 PCW), and cell counts were modelled using a negative binomial generalised model, with Benjamini-Hochberg weighted correction as previously described^9,99^, to test the effects of age (age bins) together with cell sorting correction (milopy.core.DA_nhoods). Significantly differentially abundant neighborhoods were detected by (SpatialFDR<0.1, logFC <0) for early enriched neighborhoods and (SpatialFDR<0.1, logFC >0) for late neighborhoods (**Supplementary Table 3**).

#### Cell state predictions: adult hair follicles, embryonic macrophages, blood vessel organoid

To compare prenatal skin cells with external datasets (adult hair follicle, embryonic macrophages, blood vessel organoid)^14,97,128^, the datasets were down-sampled to have roughly balanced cell counts per annotated cell type before integration with Harmony (v0.0.5)^153^, treating datasets as batches and within dataset batches as covariates (donor for prenatal skin, site for embryonic macrophages and group (cell line: day of culture) for blood vessel organoid).

Comparison of cell type correspondence between datasets and probability prediction was carried out using a LR framework similar to CellTypist package^155^. A model was built using the implementation of the linear_model.LogisticRegression module from sklearn package (v1.1.3) (parameters: penalty: L2, solver: saga, regularisation strength C=0.1) and trained on the gene expression matrix of the training dataset using all genes that passed QC. The resulting model was used to predict the labels in the target dataset. The correspondence between predicted and original labels in target dataset was computed as Jaccard index and visualised as heatmaps. For comparison of the prenatal skin macrophages with embryonic macrophages, the embryonic macrophage dataset was used as training data and prenatal skin macrophages as query; for comparison of the blood vessel organoid with prenatal skin, the prenatal skin dataset (down-sampled to max 500 cells per cell type) was used as training data and the blood vessel organoid data as query; for comparison of hair follicle data, merged prenatal and organoid data was used as training data and adult dataset as query.

#### FRZB comparison across developing organs

To compare gene expression of *FRZB* in fibroblasts across developing organs, a scRNA-seq stromal dataset from multiple developing organs^9^ (available from: https://developmental.cellatlas.io/fetal-immune) was used, which also includes our prenatal skin scRNA-seq data. The data was normalised to 1e4 counts per cell (scanpy.pp.normalize_total), log1p transformed (scanpy.pp.log1p) and subsetted to fibroblast cell types only to plot expression of *FRZB* by organ across gestation time.

#### Trajectory analysis

The CellRank package (v1.5.2) was used to define cell transition matrices, lineage drivers and rank fate probabilities of terminal state transitions across annotated lineages in prenatal skin for endothelial cells and fibroblasts and in the skin organoid for keratinocytes. Using pp.moments (n_pcs=10, n_neighbours=30) from the scVelo package (v0.3.0), first order kinetics matrices were imputed. Using the palantir kernel and the velocity kernel in CellRank, a mixed probability transition matrix was computed with each kernel weighing 50 %. Schur matrix Eigen decomposition (n_components=25, method=‘brandts’) identified macrostates, terminal stages and initial stages. Lineage drivers were then computed for each state using compute_lineage_drivers from CellRank and pseudotime and latent time computed in scVelo (**Supplementary Table 6**).

#### In vivo-in vitro trajectory alignment analysis

We used Dynamic Programming (DP) based alignment to evaluate agreement between the single-cell trajectories of prenatal skin and skin organoid fibroblasts which describe the *in vivo* and *in vitro* differentiation lineages from *HOXC5^+^* early fibroblasts to dermal papilla. Genes2Genes (G2G)^63^ is a Bayesian Information-theoretic DP framework that consistently captures matches and mismatches between two trajectories at both gene-level and cell-level. G2G outputs an optimal trajectory alignment which describes a non-linear mapping of *in vivo* and *in vitro* pseudotime points in sequential order. This is based on the cost of matching or mismatching the gene expression distributions of each pair of organoid-reference time points, computed as a statistic of entropy difference between the two hypotheses under the Minimum Message Length^156^ criterion. This statistic is a Shannon Information distance, calculated in the unit of information, *nits*. Given any gene set, G2G runs DP alignment for each gene, outputting a 5-state alignment string over matches (one-to-one/one-to-many/many-to-one) and mismatches (insertions and deletions – gaps) between the *in vivo* and *in vitro* pseudotime points in sequential order, which is analogous to a DNA/protein alignment output. It then computes an alignment similarity measure (i.e., the percentage of matches across the alignment string) for each gene (**Supplementary Table 8**). G2G also generates an aggregated cell-level alignment by averaging across all gene-level alignments, resulting in an overall alignment similarity measure as well.

Using G2G, we examined the *in vivo* reference versus *in vitro* query alignment in terms of 1363 human transcription factors^157^. These TFs were taken after filtering zero expressed genes and genes expressed in less than 10 cells. Given the reference and organoid log1p normalised gene expression matrices of cells and their pseudotime estimates computed using CellRank^158^, G2G generated fully descriptive TF-level alignments, as well as an aggregated cell-level alignment across those TF-level alignments. Prior to alignment, the optimal pseudotime binnings of reference and organoid were determined using the *OptBinning*^159^ python package, based on the given pseudotime estimates distribution, resulting in 13 optimal bins for each. Next the DP alignment was run for each TF, and the TF clusters of different alignment patterns (i.e. early mismatches, mid mismatches, late mismatches and complete mismatches) were identified using the G2G function that runs agglomerative hierarchical clustering over the TF-level alignment outputs.

#### Cell-cell interaction analysis

CellPhoneDB (v3.0.0) package^160^ was used to infer cell-cell interactions within the prenatal skin scRNA-seq dataset overall and in early/late gestation and within the skin organoid scRNA-seq dataset overall. In the overall analysis, we randomly subsampled each cell type into no more than 200 cells. The subsampled dataset was analysed using the permutation-based method to establish statistical significance (p-value cut-off = 0.05). For the analysis by early/late gestation, the prenatal skin scRNA-seq dataset was first split into early (≤ 11 PCW) and late (≥ 12 PCW) gestation datasets which were then subsampled (no more than 200 cells per cell type) and analysed individually (p-value cut-off = 0.05). A summary output file was created for each analysis run, compiling the interactions for each cell pair (p<0.05) and adjusting p-values for multiple testing (FDR set at 0.05) (**Supplementary Tables 7, 19**). Circos plots (Circlize package (v 0.4.15)^161^) were used for downstream visualisations of selected significant (adjusted p-value <0.05) interactions between co-locating cell types.

To explore inferred interactions between macrophage subsets and endothelial cells (**Extended Data Fig. 8a**), we aggregated the interactions predicted for each macrophage subset and the different subtypes of endothelial cells (early endothelial cells, arterioles, capillary arterioles, capillaries, postcapillary venules, venules) by averaging the means and using the minimum of the adjusted p-values as previously described^9^. A curated list of aggregated interactions were plotted for visualisation using ggplot2 (v 3.3.6). A similar approach was adopted for assessing interactions between hair follicle dermal and epidermal cells in prenatal skin: for each subset of hair follicle dermal cells, the interactions with early epithelial cells (≤11 PCW; Immature basal) or late epithelial cells (≥12 PCW; *DPYSL2^+^* basal, *POSTN^+^* basal, Placode/matrix, Outer root sheath, Companion layer, Inner root sheath, Cuticle/Cortex) were aggregated and the top 10 interactions per cell pair visualised using heatmap (**Fig. 2i**). The same analysis was performed to obtain the top 10 interactions in skin organoid hair follicles (**Extended Data Fig. 4g**), defining early/late to match corresponding cell states as in prenatal skin. The top 10 interactions identified in prenatal skin hair follicles were also plotted within the skin organoid data to highlight similarities and differences between the two (**Fig. 2i**).

#### Comparison with adult fibroblasts

scRNA-seq data from prenatal skin and adult healthy skin (with original annotations)^13^ were pooled, retaining only genes expressed in at least 1 cell for each dataset, and subsetted to the cell group of interest (fibroblasts). Differentially expressed genes between the adult and prenatal skin fibroblasts were derived using the Wilcoxon rank-sum test implementation in scanpy and adjusted for multiple testing using the Benjamini–Hochberg method (scanpy.tl.rank_genes_groups, method = “wilcoxon”, corr_method=”benjamini-hochberg”). A selected list of genes was plotted to highlight differences between prenatal and adult skin fibroblasts.

#### Gene set enrichment analysis

Gene set enrichment analysis was performed using the implementation of the Enrichr workflow^122^ in the python package GSEApy (https://gseapy.readthedocs.io/) with Gene Ontology (GO) Biological Process (2021) and Molecular Signatures Database (MSigDB) Hallmark (2020) as query databases. To determine the significantly overexpressed genes for gene set enrichment analysis, we first identified the differentially expressed genes between cell types for each cell group of interest (macrophages) using the Wilcoxon rank-sum test implementation in Scanpy (scanpy.tl.rank_genes_groups, method = “wilcoxon”). Genes with differential expression logFC > 2 and adjusted p-value < 0.01 were considered as significantly overexpressed.

For comparison between early and late cell states, for cell types of interest (*WNT2*^+^ fibroblast), we first identified the index cells belonging to early neighbourhoods (SpatialFDR < 0.1, logFC < 0) and late neighbourhoods (SpatialFDR < 0.1, logFC > 0) based on Milo^18^ differential abundance testing as described above (**Supplementary Table 3**). Differentially expressed genes between early and late cell states were computed using the Wilcoxon rank-sum test implementation in scanpy (scanpy.tl.rank_genes_groups, method = “wilcoxon”). Genes expressed in less than around 10% cells were excluded. Genes with differential expression logFC > 1 and adjusted p-value < 0.01 were considered as significantly overexpressed for gene set enrichment analysis using GSEApy (https://gseapy.readthedocs.io/), with Gene Ontology (GO) Biological Process (2021) as query database. The list of significantly over-expressed genes for each cell type/state where differential expression testing and gene set enrichment analysis were carried out can be found in **Supplementary Tables 11, 12**.

#### Gene module scoring

Gene module scoring was performed using the sc.tl.score_genes function in scanpy. For angiogenesis gene modules, pre-defined gene sets from the Gene Ontology Biological Process Database (2021) in Enrichr libraries^123^ were used (downloaded from: https://maayanlab.cloud/Enrichr/#libraries). For endothelial cell modules, gene sets defining tip, stalk, arteriole, venule, lymphatic, capillary and hypoxia scores (**Extended data Fig. 9g, h**) were derived from published literature^125–127^. The list of genes for each gene module is provided in **Supplementary Tables 18, 20**. The score for each module is the average expression of the gene set provided subtracted with the average expression of a reference set of genes. The reference set comprised 100 genes (ctrl_size=100) which were randomly sampled from all genes in the dataset (default gene_pool) with 25 expression level bins (n_bins=25) used for sampling. For angiogenesis modules, the mean module scores were computed for each cell type of interest (e.g., *LYVE1*^+^ macrophage) and z-score normalised for visualisation.

#### Gene regulatory network analysis

The PySCENIC package (v0.11.2) and pipeline were used to identify transcription factors and their target genes in the combined prenatal skin and skin organoid scRNA-seq datasets. The ranking database (hg38 refseq-r80 500bp_up_and_100bp_down_tss.mc9nr.feather), motif annotation database (motifs-v9-nr.hgnc-m0.001-o0.0.tbl) were downloaded from the Aert’s laboratory github page. The tool was run 10 times, with a data set comprising at most 200 cells per cell type x tissue pair (where tissue is prenatal skin or skin organoid). For each run an adjacency matrix of transcription factors and their targets was generated and pruned using the Aert’s group suggested parameters. Only regulons present in at least 8 out of 10 runs were used in the analysis. PySCENIC was used to calculate the Regulon Specificity Score for each cell type x tissue pair using *aucell* function. An average was computed over the multiple runs. These average scores were used to compare regulon activity between prenatal skin and skin organoid. A gene interaction network was first built by querying the STRING database with GATA2 target genes, then pruned to only keep genes reported as associated with GATA2. The list was further truncated to 12 genes, by keeping genes that were 1) transcription factors in the five most active regulons detected in fetal skin and/or 2) organoid capillary arterioles, and/or 3) associated with pseudotime (i.e., in trajectories) and/or 4) VEGF receptors and/or 5) in the selected GO terms chose for their role in angiogenesis, extracellular matrix organization, or cell migration, communication, proliferation, or death (’GO:0045765’, ‘GO:0001568’, ‘GO:0030334’, ‘GO:0010646’, ‘GO:0001936’, ‘GO:0045446’, ‘GO:0002040’, ‘GO:0030155’, ‘GO:0010941’, ‘GO:0030198’).

#### Comparison of pro- and anti-angiogenic genes between prenatal skin and skin organoid

The prenatal skin and skin organoid data sets were integrated with Harmony (v0.0.5)^153^ as described above. Differential expression analysis was performed between prenatal skin and skin organoid cells (all cell types) using the standard scanpy workflow (scanpy.tl.rank_genes_groups, method = “wilcoxon”). Identified DEGs were filtered to only retain those coding for secreted proteins (**Supplementary Table 29**)^162^. Gene set enrichment analysis was performed on downregulated and upregulated genes separately, using the implementation of the Enrichr workflow^122^ in the python package GSEApy (https://gseapy.readthedocs.io/) with Gene Ontology (GO) Biological Process (2021) as query database. Significant GO terms (adjusted p-value < 0.05) (**Supplementary Tables 24, 25**) were filtered based on their relevance to vasculature. Only DEGs involved in pathways thereby selected were chosen and their role in prenatal skin angiogenesis checked in the literature.

#### NicheNet analysis

We used NicheNet^136^ (v.1.1.1) to infer ligand-target gene links by combining scRNA-seq data (prenatal skin and skin organoid) of interacting cells (sender and receiver cells) with existing knowledge on signalling and gene regulatory networks. An open-source R implementation including integrated data sources used in the analysis are available at GitHub (https://github.com/saeyslab/nichenetr). Nichenet’s ligand-activity analysis first assesses and ranks ligands in the sender cell type (macrophage subsets) which best predict observed changes in expression of target genes of interest in the receiver cell types (endothelial cells) compared to background genes. Potential ligands were defined as all ligands in the NicheNet model which were expressed in at least 10% of cells in each macrophage (sender) cluster and which had at least one specific receptor expressed in at least 10% of endothelial (receiver) cells. Target genes of interest were identified as differentially expressed genes between conditions (prenatal skin vs skin organoid) in receiver cells using FindMarkers function in NicheNet (adjusted p-value ≤0.05 and average log2 fold change >0.25, expressed in at least 10% of endothelial cells). Background genes were all genes in the NicheNet model which were expressed in at least 10% of receiver cells.

Ligands were prioritised based on ligand activity scores, calculated as the Pearson correlation coefficient between a ligand’s target predictions and the observed target gene expression (**Supplementary Table 26**). The top 20 ligands were used to predict active target genes (top 200 overall) and construct the active ligand-target links (**Supplementary Table 27**). Receptors of the top-ranked ligands were identified from the NicheNet model, filtering for only bona-fide ligand-receptor interactions documented in the literature and publicly available databases (**Supplementary Table 28**). To infer signalling paths, we defined our ligand (*VEGFA*, in red) and target genes (*GATA2*, in blue) of interest. NicheNet identifies which transcription factors best regulate the target genes and are most closely downstream of the ligand based on weights of the edges in its integrated ligand-signalling and gene regulatory networks. The shortest paths between these transcription factors and the defined ligand are selected and genes along these paths are considered as relevant signalling mediators (in grey). Signalling mediators are prioritised by cross-checking the interactions between the defined ligand, target genes and identified transcription factors and signalling mediators across the different integrated data sources in NicheNet.

#### Spatial data analysis

Spatial transcriptomics data was mapped using Space Ranger v.2.0.1 using GRCh38-1.2.0 reference. In parallel, we manually selected skin-overlapping spots in embryonic limb data^16^, comprising samples from the following ages: 6 PCW (n=2, replicate=2 each) and 8 PCW (n=1, replicate=3). To map cell types identified by scRNA-seq in the profiled spatial transcriptomics slides, we used the Cell2location (v0.1) method^19^. Firstly, we trained a negative binomial regression model to estimate reference transcriptomic profiles for all the cell types profiled with scRNA-seq in the organ. We excluded very lowly expressed genes using the filtering strategy recommended by Cell2location authors (cell_count_cutoff=5, cell_percentage_cutoff2=0.03, nonz_mean_cutoff=1.12). Cell types where less than 20 cells were identified in samples ≤10 PCW were excluded from the reference. Individual 10x samples were considered as a batch, donor and chemistry type information was included as categorical covariate. Training was performed for 250 epochs and reached convergence according to manual inspection. Next, we estimated the abundance of cell types in the spatial transcriptomics slides using reference transcriptomic profiles of different cell types. All slides were analysed jointly. The following Cell2location hyperparameters were used: (1) expected cell abundance (N_cells_per_location) = 30; (2) regularisation strength of detection efficiency effect (detection_alpha) = 20. The training was stopped after 50,000 iterations. All other parameters were used at default settings. Cell2location estimates the posterior distribution of cell abundance of every cell type in every spot. Posterior distribution was summarised as 5% quantile, representing the value of cell abundance that the model has high confidence in, and thus incorporating the uncertainty in the estimate into values reported in the paper and used for downstream co-location analysis.

To identify microenvironments of co-locating cell types, we used non-negative matrix factorisation (NMF). We first normalised the matrix of estimated cell type abundances by dividing it by per-spot total abundances. Resulting matrix Xn of dimensions *n* × *c*, where n is the total number of spots in the Visium slides and *c* is the number of cell types in the reference was decomposed as *X*n = *WZ*, where W is a *n* × *d* matrix of latent factor values for each spot and Z is a *d* × *c* matrix representing the fraction of abundance of each cell type attributed to each latent factor. Here latent factors correspond to tissue microenvironments defined by a set of co-localised cell types. We use the NMF package for R^163^, setting the number of factors d = 10 and using the default algorithm^164^. NMF coefficients were normalised by a per-factor maximum. We ran NMF 100 times and constructed the coincidence matrix. Then we selected the best run based on the lower mean silhouette score calculated on the coincidence matrix. If more than one run had the minimal mean silhouette, we selected one with smaller deviance (as reported by NMF function).

For cell type abundance correlation analysis, we used a per-spot normalised Xn matrix. Pearson correlation coefficient was calculated for each pair of cell types (all possible pairs computed) and each sample. For visualisation of correlation analysis, selected cell pairs were plotted, guided by NMF results and which cell groups/categories formed microenvironments, e.g., macrophages formed microenvironments with endothelial cells (microenvironments 1 and 5), with neuronal cells (microenvironments 1 and 5) and fibroblasts (microenvironments 1, 4 and 5) in **Fig. 1d**.

## Acknowledgements

We thank Aidan Maartens for scientific writing support. We acknowledge funding from the Wellcome Human Developmental Biology Initiative (WT215116/Z/18/Z). M.H. is funded by Wellcome (WT107931/Z/15/Z), The Lister Institute for Preventive Medicine and NIHR and Newcastle Biomedical Research Centre. S.A.T. is funded by Wellcome (WT206194) and the ERC Consolidator Grant DEFINE. L.J. is funded by a Newcastle Health Innovation Partners Lectureship. N.H.G. is funded by an MRC Clinical Research Training Fellowship (MR/W015625/1). B.O. is funded by a Wellcome 4Ward North Clinical Training Fellowship. Some of this research was supported by the NIHR Cambridge Biomedical Research Centre (NIHR203312); the views expressed are those of the authors and not necessarily those of the NIHR or the Department of Health and Social Care and Wellcome (203151/Z/16/Z). This publication is part of the Human Cell Atlas – www.humancellatlas.org/publications/[humancellatlas.org].

## Author contributions

Conceptualization: M.H., S.A.T., and K.K. Funding acquisition: M.H., S.A.T. Supervision: M.H. and L.G. Data curation: N.H.G., N.H., B.O., P.M., S.B., D.H., D.B.L. and P.Ma. Formal analysis: N.H.G., N.H., B.O., C.A., R.A.B., A.R.F., E.W., D.S., W.M.T., P.M., I.G., A.R., D.M.P., S.B. and P.Ma. Software: D.H., D.B.L., T.L., J.M., O.B. Investigation: C.A., R.A.B., A.R.F., F.T., M.M., E.S., B.R., K.R., K.S., J.E., V.R., J.F., D.M.P., E.P, J-E.P., S.P., I.M. and L.G. Methodology: N.H., E.W., D.S., I.G., C.A., R.A.B, E.S. and P.Ma. Resources: P.H., S.L., I.G., V.R., A.Fi., S.Li., R.B., R.V.T., A.P.L., S.S., J.K., C.D., J.L., N.R., M.N., B.T. and K.K. Writing - original draft: N.H.G., N.H., B.O., C.A., A.R.F., E.W., D.S., B.R., S.B., P.Ma., L.G., M.H. Writing - review and editing: N.H.G., N.H., B.O., I.G., E.S., L.J., G.R., C.G., E.P., A.D., A.P.L., S.S., J.L., N.R., E.O’T., M.K., L.G., K.K., S.A.T. and M.H. Visualization: N.H.G., B.O., C.A., M.M., K.S., D.B.L., D.H., T.L., J.M., O.B., L.G. and M.H.

## Additional information statement

For additional information regarding reprints and permissions, correspondence and requests for materials must be addressed to Muzlifah Haniffa at.

## Data availability statement

The datasets generated and/or analysed during the current study are available in the following repositories: Prenatal scRNA-seq skin data is available on ArrayExpress under E-MTAB-11343, E-MTAB-7407 and E-MTAB-13071 accessions. Prenatal skin TCR-seq data is available under E-MTAB-13065 accession. Skin organoid scRNA-seq data is available on GEO under GSE147206, GSE188936, and GSE231607 accessions. Visium limb data is available under E-MTAB-10367 and Visium facial and abdominal data are deposited in ArrayExpress under E-MTAB-13024. Embryonic macrophage scRNA-seq data is available on GEO under accession numbers GSE13345 and GSE137010. All of the blood vessel organoid scRNA-seq data analysed as part of this study are included in the pre-print article from Nikolova et al^128^. Adult healthy skin scRNA-seq is available on ArrayExpress under E-MTAB-8142. Adult hair follicle scRNA-seq data is accessible from GEO under GSE129611. Processed data can be accessed on our web-portal https://developmental.cellatlas.io/fetal-skin (password: fs2023).

## Code availability statement

Single-cell and spatial data were processed and analysed using publicly available software packages. Python/R code and notebooks for reproducing these analyses are publicly available at https://doi.org/10.5281/zenodo.8164271.

---

**Extended Data Fig. 1.**
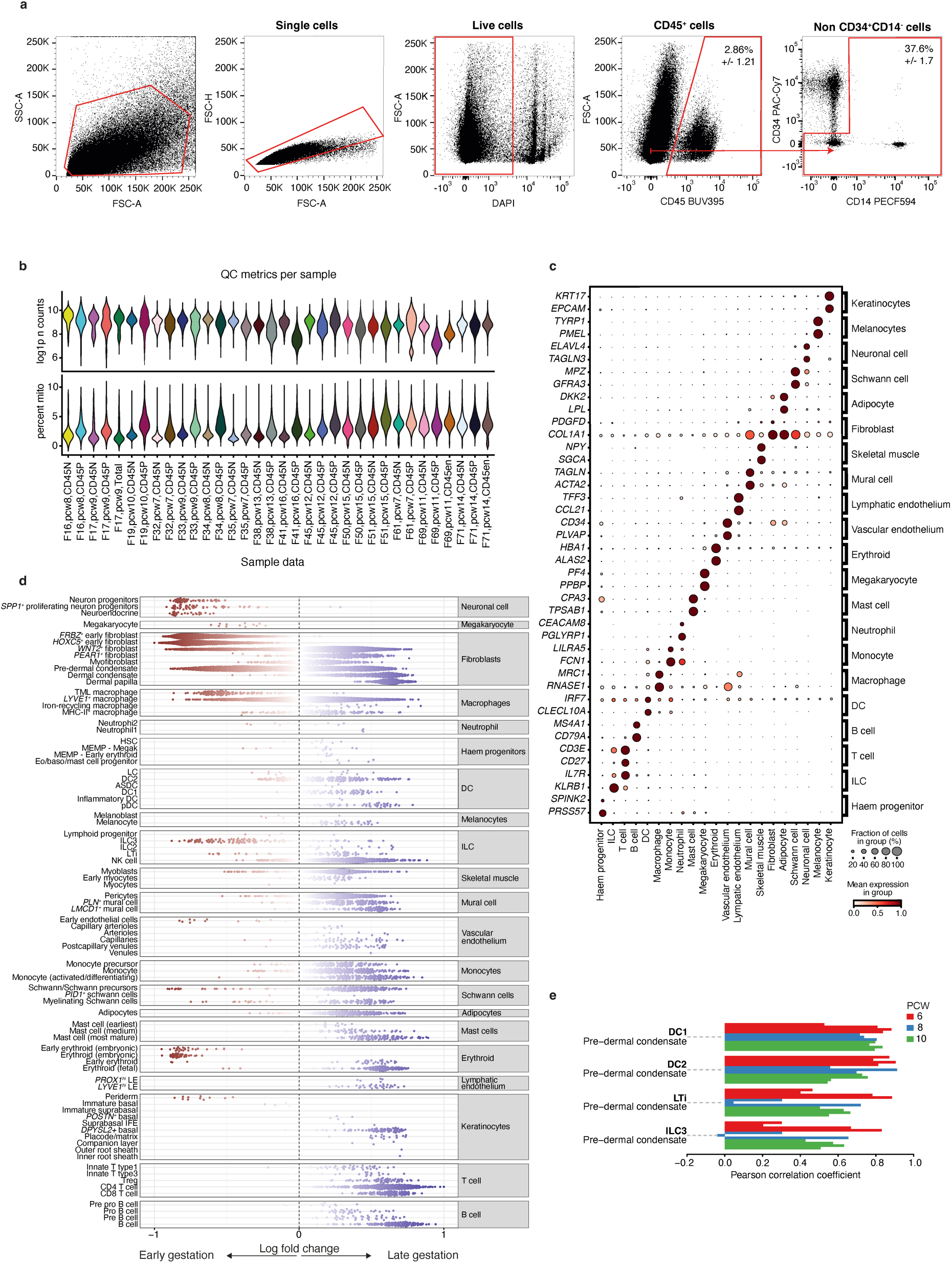
Temporal and spatial composition of human prenatal skin. **(a)** Prenatal skin cells isolation by fluorescence-activated cell sorting into CD45^+^ and CD45^-^ fractions (n=15); from the CD45^-^ fraction we further isolated all cells that were not within the CD34^+^CD14^-^ gate (n=2) to enrich for endothelial cells and keratinocytes. Representative data from n=1 is shown as mean percentage +/- SD values. **(b)** Quality control plots showing frequency distribution of UMI counts (log1p-transformed) and percent of UMI counts in mitochondrial genes per sample fraction. **(c)** Dot plot showing variance-scaled, mean expression (dot colour) and percent of expressing cells (dot size) of defining genes for cell states corresponding to Fig. 1b. **(d)** Milo beeswarm plot showing differential abundance of neighbourhoods in prenatal skin across gestation time, annotated by refined cell labels. Red/blue neighbourhoods are significantly enriched in earlier/later gestation respectively. Colour intensity denotes degree of significance. **(e)** Bar plot showing cell type co-location, indicated by positive Pearson correlation coefficient, for selected cell type pairs (pre-dermal condensate and immune cells: DC1, DC2, LTi and ILC3). Pearson correlation coefficients were calculated across all skin-covered spots of Visium samples; each sample is shown by an individual bar. ASDC: *Axl+Siglec6+* dendritic cells; DC: dendritic cells; HSC: hematopoietic stem cells, ILC: innate lymphoid cells, LC: Langerhans cells, LTi: lymphoid tissue inducer cells, CD45en: CD45 negative fraction enriched for keratinocyte/endothelial cells; CD45N: CD45 negative; CD45P: CD45 positive; pDC: plasmacytoid dendritic cells, TML macrophage: *TREM2^+^* microglia-like macrophage.

**Extended Data Fig. 2.**
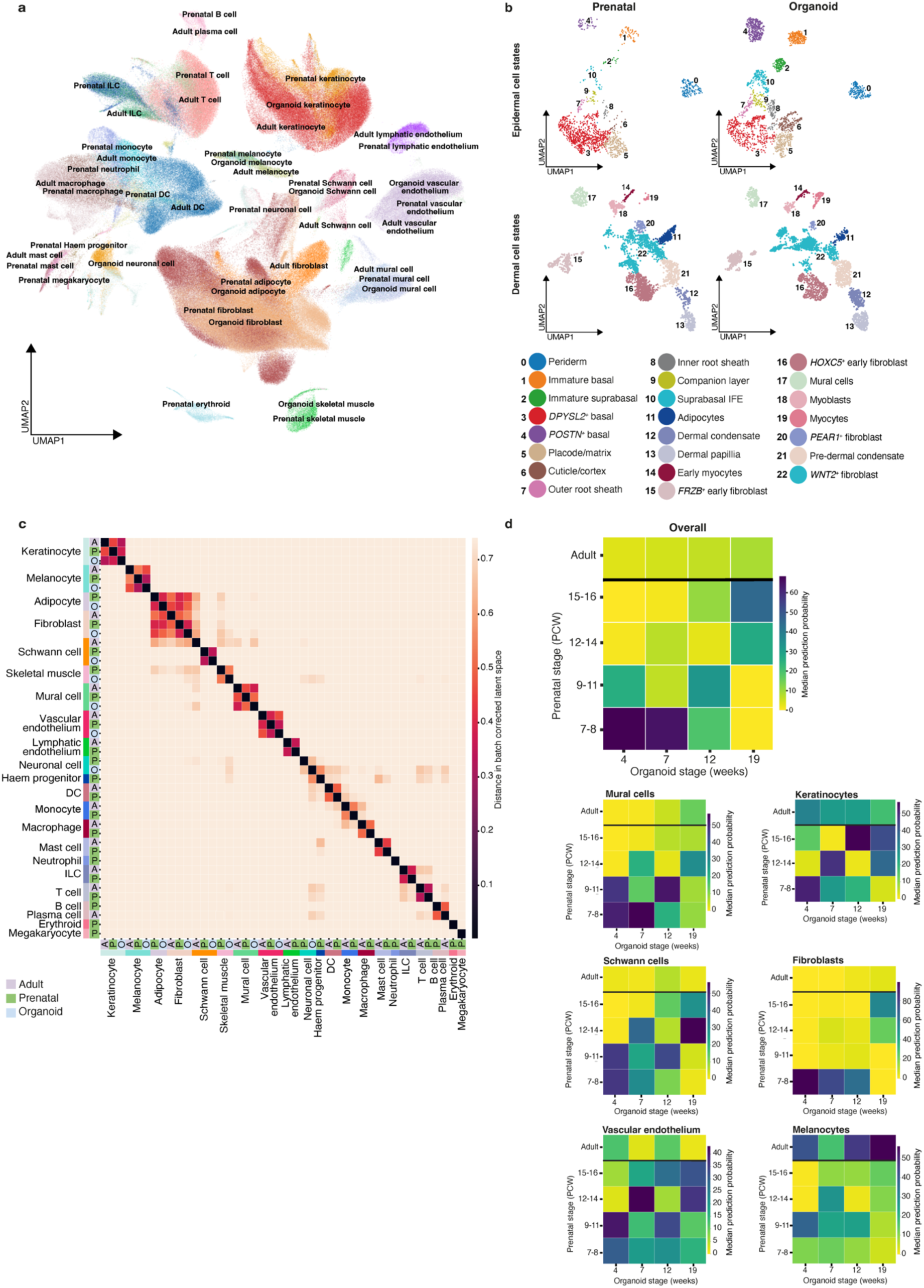
Comparison of the skin organoid with prenatal and adult skin. **(a)** UMAP visualisation of the integrated prenatal skin, adult skin^13^ and skin organoid^15^ scRNA-seq datasets, coloured by broad cell types. **(b)** UMAP visualisations of integrated data from prenatal skin and skin organoid, coloured by epidermal (top) and dermal (bottom) cell types in prenatal skin (left) and skin organoid (right). **(c)** Heatmap showing conserved cell states (measured by distance in principal component space) between prenatal skin, adult skin^13^ and skin organoid^15^ for broad cell categories. **(d)** Heatmap showing prediction probabilities (overall and per broad cell category) for a logistic regression model trained on time-encoded prenatal skin and adult skin data (y-axis)^13^ and projected onto time-encoded skin organoid data^15^ (x-axis). Colour scale indicates median prediction probabilities.

**Extended Data Fig. 3.**
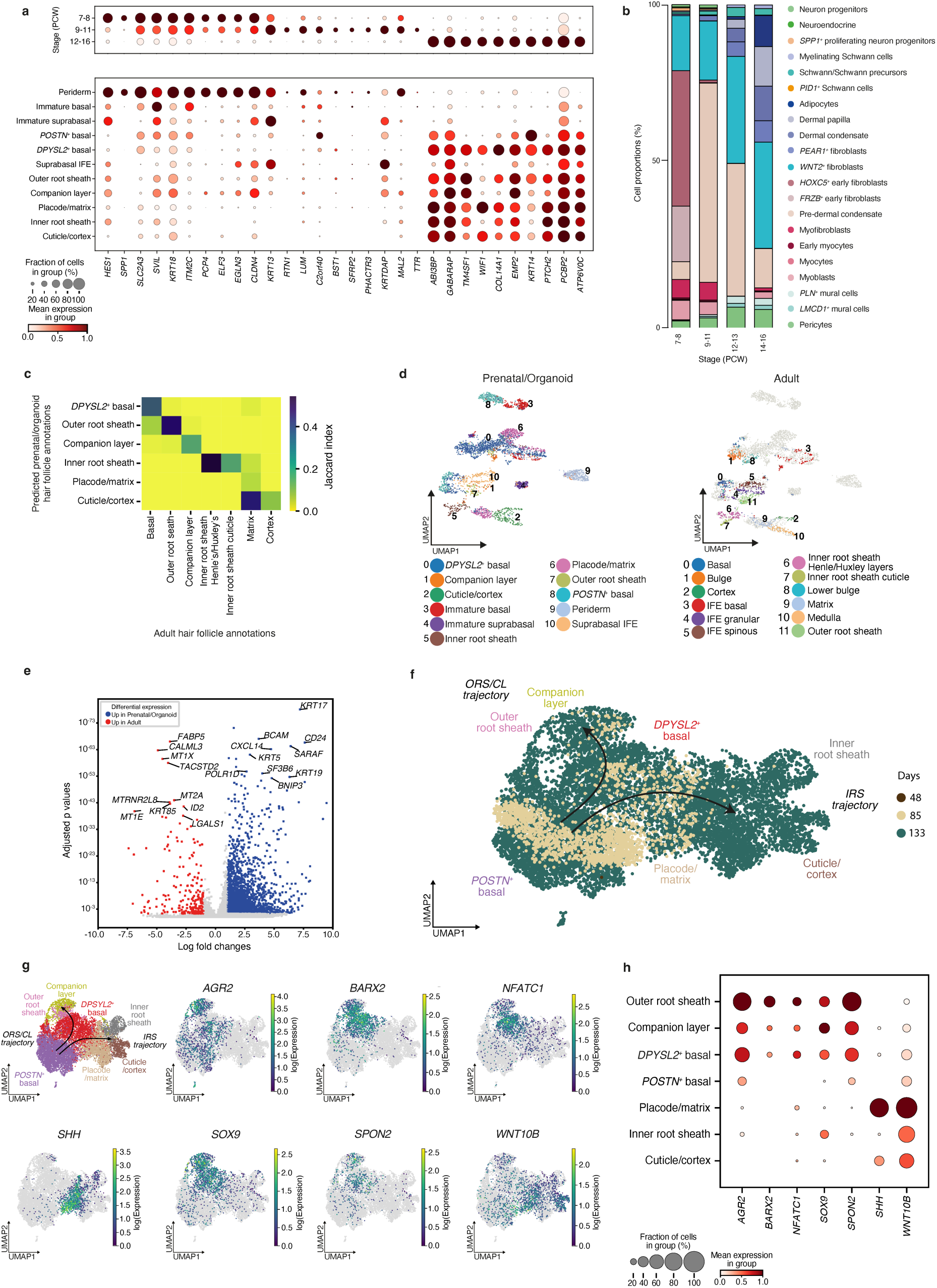
Differentiation of prenatal hair follicle epidermal cells and comparison with adult hair follicles. **(a)** Dot plot showing variance-scaled, mean expression (dot colour) and percent of expressing cells (dot size) of DEGs between gestational stage groups (grouped PCW) (top) and expression of the same genes by different epidermal cell states (bottom). **(b)** Bar plot showing the proportions of stromal cell states across gestational age in prenatal skin. Bar colours represent cell states. **(c)** Heatmap showing the correspondence (measured by Jaccard index) between prenatal skin/skin organoid (y-axis) and adult (x-axis) epidermal and hair follicle cell states for a logistic regression model trained on adult hair follicle data^14^ and projected onto integrated prenatal skin/skin organoid data. **(d)** UMAPs showing clustered cell states in integrated data from adult hair follicles^14^ and prenatal/skin organoid, coloured by prenatal skin/skin organoid cell types (left) and adult cell types (right). **(e)** Volcano plot showing differentially expressed genes between prenatal/organoid placode/matrix cells and adult matrix cells^14^. **(f)** Inferred pseudotime trajectory of skin organoid epidermal cell states differentiating along the ‘ORS/CL’ and ‘IRS’ trajectories, coloured by days of culture. **(g)** Inferred pseudotime trajectory of skin organoid epidermal cell states differentiating along the ‘ORS/CL’ and ‘IRS’ trajectories, coloured by gene expression (log-transformed). **(h)** Dot plot showing variance-scaled, mean expression (dot colour) and percent of expressing cells (dot size) in prenatal skin of genes expressed along the ‘ORS/CL’ and ‘IRS’ trajectories. CL: companion layer, IRS: inner root sheath ORS: outer root sheath.

**Extended Data Fig. 4.**
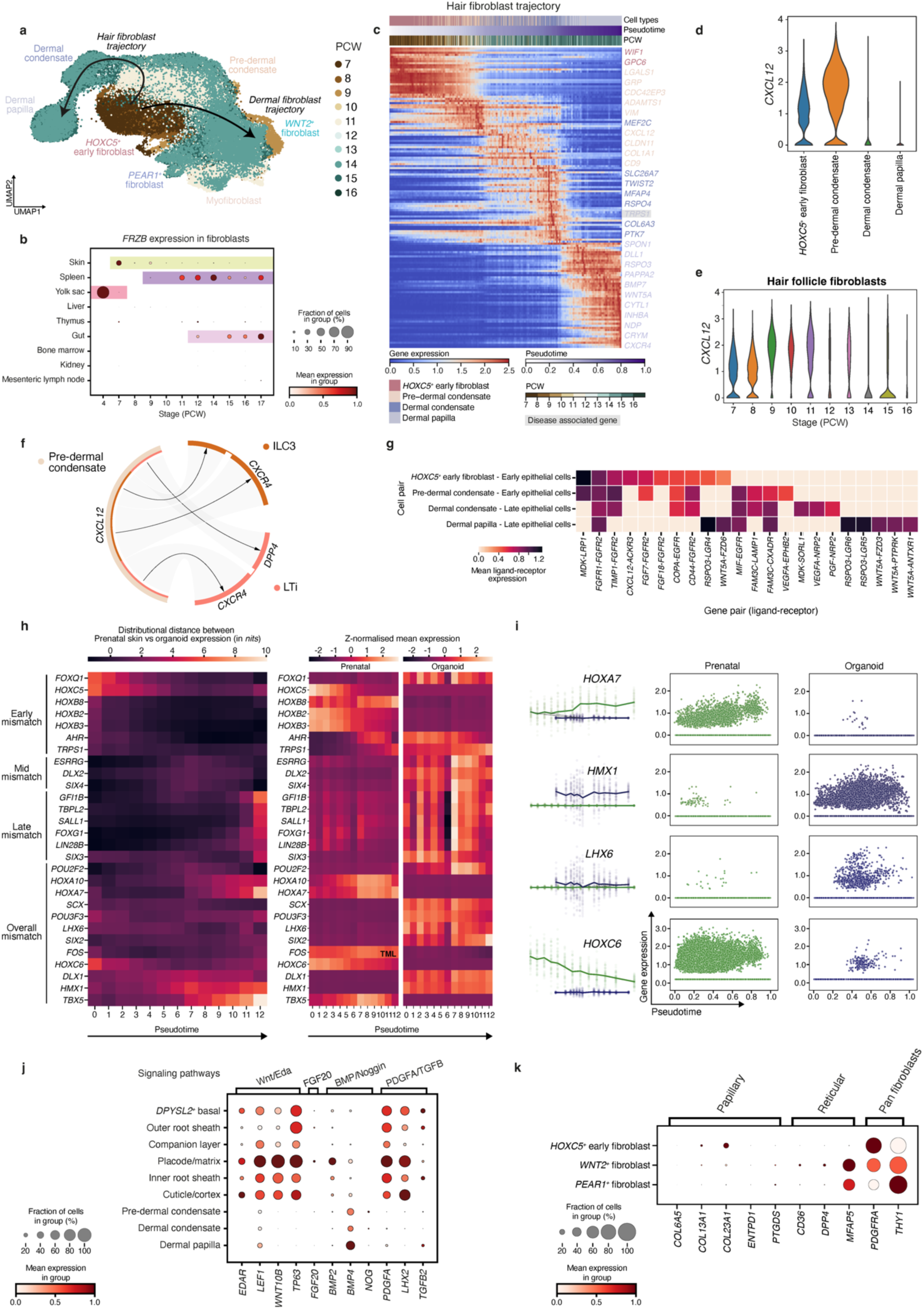
Differentiation of the prenatal hair follicle mesenchyme. **(a)** Inferred pseudotime trajectory of prenatal skin fibroblasts differentiating along the ‘hair’ and ‘dermal’ trajectories, coloured by gestational age (PCW). **(b)** Dot plot showing variance-scaled, mean expression (dot colour) and percent of expressing cells (dot size) of *FRZB* gene in fibroblasts from developing organs. Gestational ages during which individual organs are present are highlighted. **(c)** Heatmap showing differentially expressed genes across pseudotime along the ‘hair fibroblast trajectory’. Gene associated with genetic hair disorders is highlighted in grey. **(d)** Violin plot showing expression of *CXCL12* in hair mesenchymal cells (violin width proportional to counts). **(e)** Violin plot showing expression of *CXCL12* in hair mesenchymal cells by gestation (PCW) (violin width proportional to counts). **(f)** Circos plot showing selected significant (adjusted p-value<0.05) predicted interactions between pre-dermal condensate and ILC3 and LTi cells in prenatal skin. Arrows represent directionality of interactions (ligand to receptor); connection width is proportional to the CellphoneDB mean value for each ligand-receptor pair. **(g)** Heatmap showing significant (adjusted p-value <0.05) predicted interactions between hair mesenchymal cells and epithelial cells (early: Immature basal; late: *DPYSL2*^+^ basal, *POSTN^+^* basal, Placode/matrix, ORS, CL, IRS, Cuticle/cortex) in skin organoid. Top 10 interactions per cell pair are shown. Colour scale represents the mean expression values of each ligand-receptor pair in corresponding cell pairs. **(h)** Left: Heatmap showing the distributional distance (measure of Shannon information (unit: nits)) of gene expression between prenatal skin (reference) and skin organoid, as a measure of dissimilarity (mismatch) for selected, differentially expressed genes across pseudotime. Heatmap of the smoothened (interpolated) and z-normalised mean expression of the selected genes across pseudotime in prenatal skin (middle) and skin organoid (right). **(i)** Gene expression plots for representative genes in prenatal skin (green) and skin organoid (blue) across pseudotime. Left column: the interpolated log1p normalised expression (y-axis) against pseudotime (x-axis). The lines represent mean expression trends; the data points are 50 random samples from the estimated expression distribution at each time point. Right two columns: actual log1p normalised expression (y-axis) against pseudotime (x-axis) where each point represents a cell. **(j)** Dot plot showing variance-scaled, mean expression (dot colour) and percent of expressing cells (dot size) of known genes involved in hair formation^21^. **(k)** Dot plot showing variance-scaled, mean expression (dot colour) and percent of expressing cells (dot size) of fibroblast marker genes. CL: companion layer, ILC: innate lymphoid cells, IRS: inner root sheath, LTi: lymphoid tissue inducer cells ORS: outer root sheath.

**Extended Data Fig. 5.**
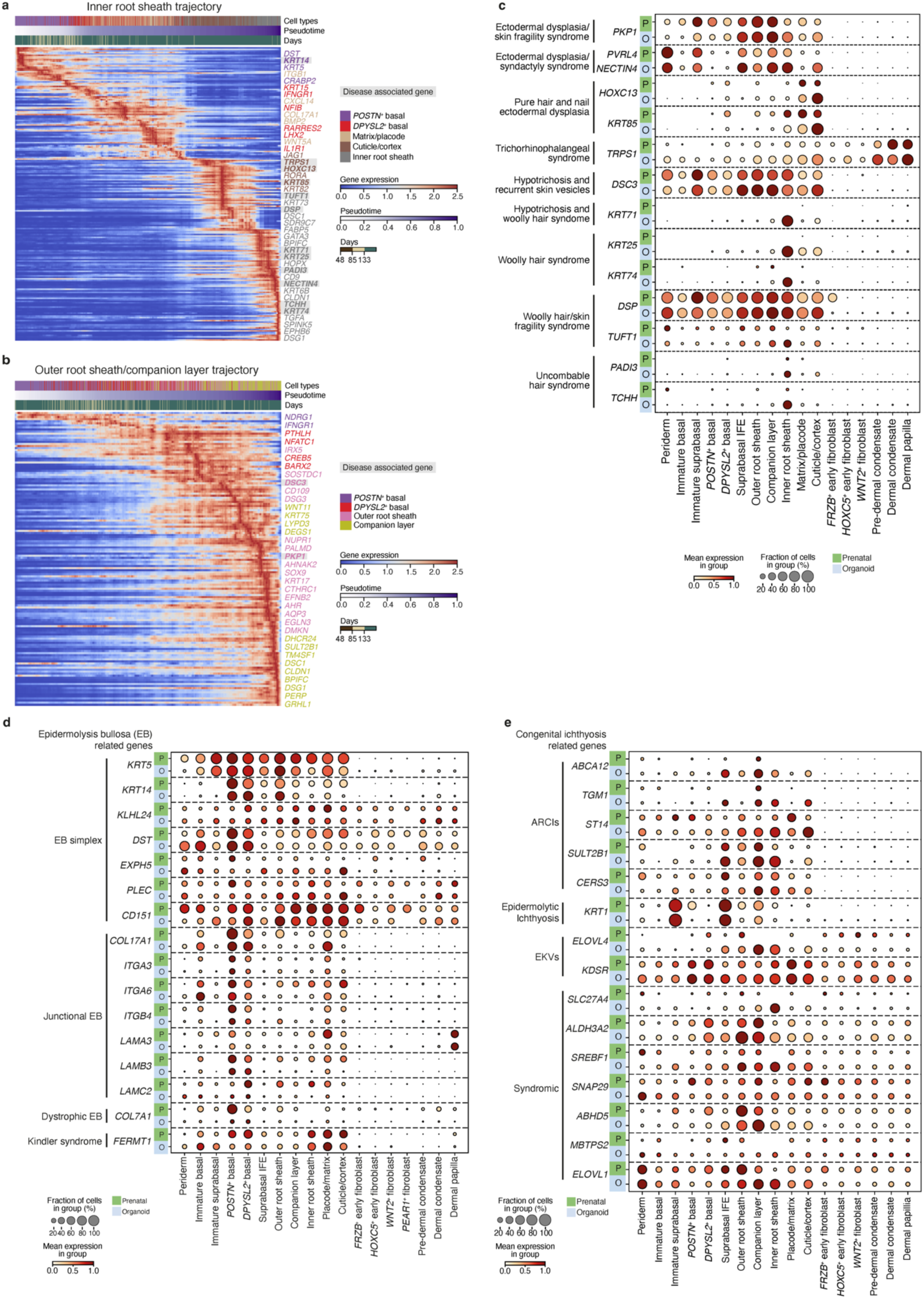
Genetic hair and skin disorders. **(a)** Heat map showing differentially expressed genes across pseudotime along the ‘Inner root sheath trajectory’. Genes associated with genetic hair disorders are highlighted in grey. **(b)** Heat map showing differentially expressed genes across pseudotime along the ‘Outer root sheath/ Companion layer trajectory’. Genes associated with genetic hair disorders are highlighted in grey. **(c)** Dot plot showing variance-scaled, mean expression (dot colour) and percent of expressing cells (dot size) of genes implicated in genetic hair diseases in prenatal skin and skin organoid skin^15^. **(d)** Dot plot showing variance-scaled mean expression (dot colour) and percent of expressing cells (dot size) of genes causing Epidermolysis Bullosa in prenatal skin and skin organoid skin^15^. **(e)** Dot plot showing variance-scaled, mean expression (dot colour) and percent of expressing cells (dot size) of genes causing congenital ichthyoses in prenatal skin and skin organoid skin^15^.

**Extended Data Fig. 6.**
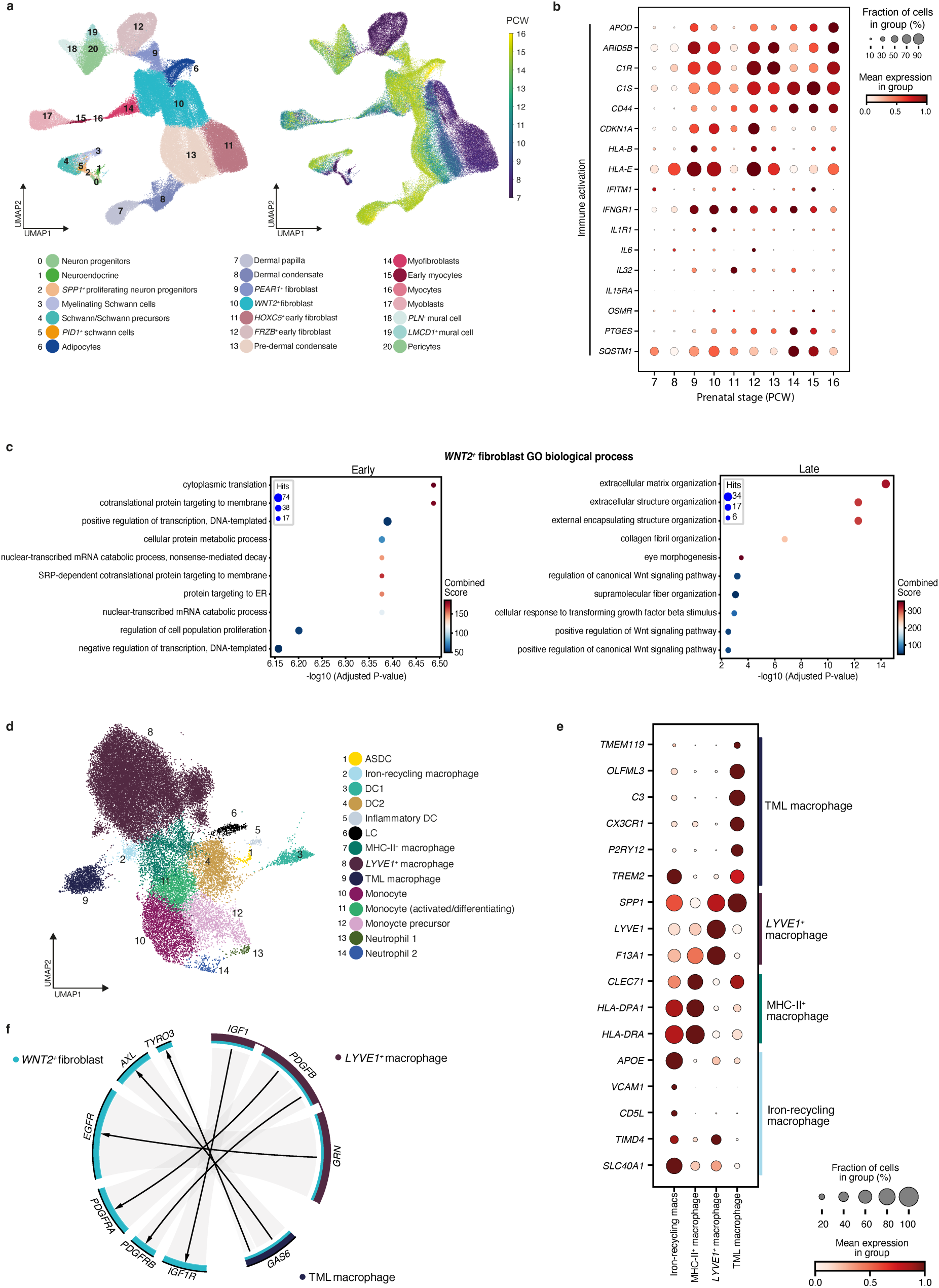
The role of early dermal fibroblasts in prenatal skin. **(a)** UMAP visualisation showing stromal cells found in prenatal skin, coloured by cell state (left) and by gestational age (PCW) right. **(b)** Dot plot showing variance-scaled, mean expression (dot colour) and percent of expressing cells (dot size) of ‘immune activation’ genes (as shown in Fig. 3a) in prenatal skin fibroblasts by gestational age (PCW). **(c)** Gene set enrichment analysis results for differentially expressed genes in Milo-defined early- and late-specific neighbourhoods of *WNT2^+^*fibroblasts. Each plot shows the top 10 enriched gene sets (using Gene Ontology Biological Process 2021). The x-axis shows the negative log_10_ of the p-value adjusted for multiple testing (Benjamini-Hochberg correction); dot size is proportional to the number of genes associated with the gene set and colour represents the combined Enrichr score calculated within GSEApy. **(d)** UMAP visualisation of the myeloid cells in prenatal skin data, coloured by cell state. **(e)** Dot plot showing variance-scaled, mean expression (dot colour) and percent of expressing cells (dot size) of marker genes^9^ used to annotate macrophage subsets in prenatal skin. **(f)** Circos plot visualisation of representative significant (adjusted p-value <0.05) predicted interactions between macrophages (*LYVE1^+^* and TMLM) and co-localising *WNT2^+^*fibroblasts in prenatal skin. Arrows represent directionality of interactions (ligand to receptor); connection width is proportional to the CellphoneDB mean value for each ligand-receptor pair. ASDC: *Axl+Siglec6+* dendritic cells; DC: dendritic cells; LC: Langerhans cells, TML macrophage: *TREM2^+^* microglia-like macrophage.

**Extended Data Fig. 7.**
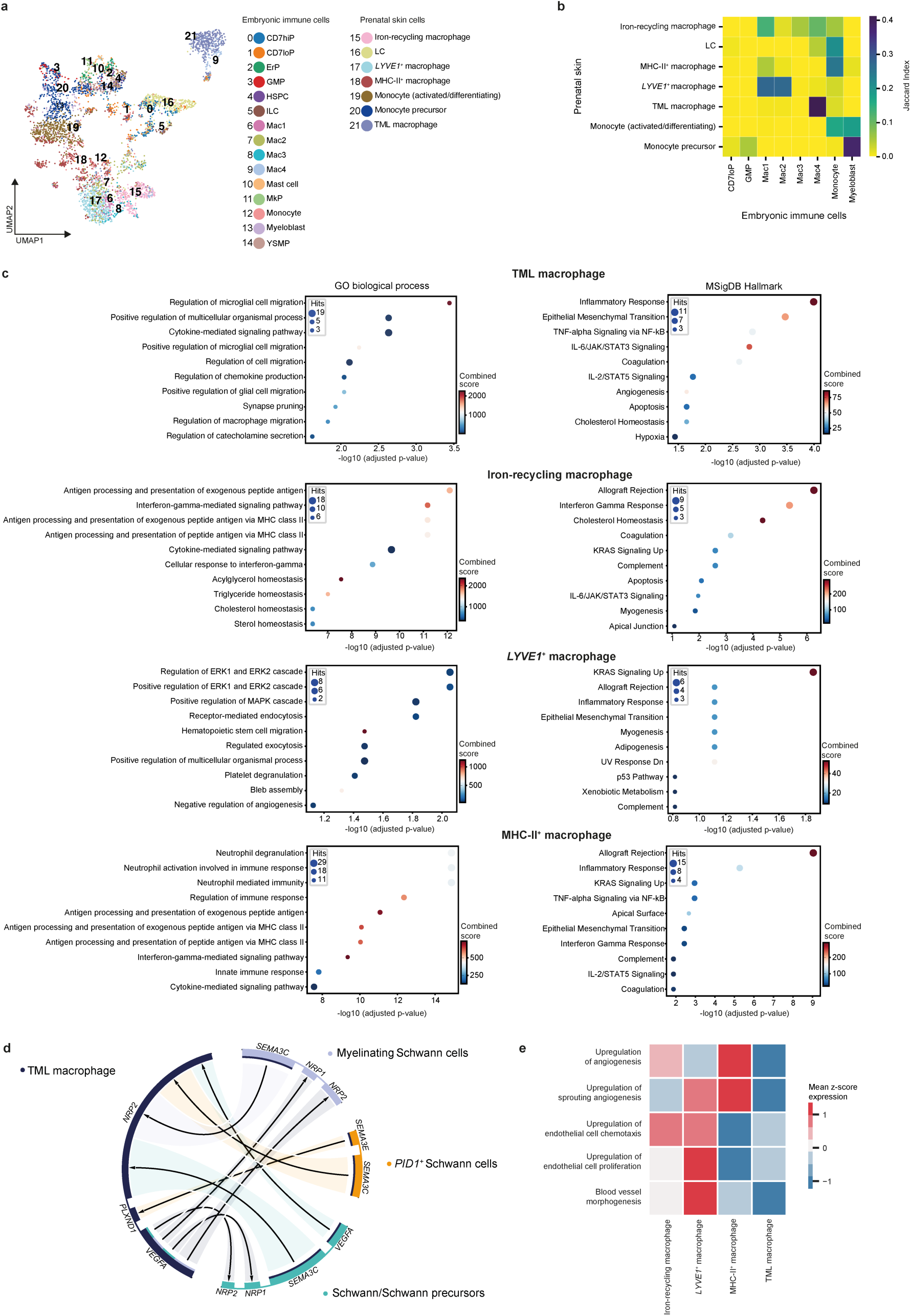
The role of macrophages in prenatal skin neurovascular development. **(a)** UMAP showing clustered cell states in integrated data from embryonic immune cells^97^ and prenatal skin myeloid cell subset. **(b)** Heatmap showing the correspondence (measured by Jaccard index) between embryonic immune cells (x-axis) and prenatal skin (y-axis) myeloid cell states for a logistic regression model trained on embryonic data^97^ and projected onto prenatal skin myeloid cell subset. TML macrophage had the highest proportion prediction to Mac4 (embryonic brain microglia). **(c)** Gene set enrichment analysis results of over-expressed genes in macrophage subsets (Iron-recycling, *LYVE1^+^*, MHC-II*^+^* and TMLM). Each plot shows the top 10 enriched gene sets (using Gene Ontology Biological Process 2021 (left) and MSigDB Hallmark 2020 (right) databases). The x-axis shows the negative log_10_ of the p-value adjusted for multiple testing (Benjamini-Hochberg correction); dot size is proportional to the number of genes associated with the gene set and colour represents the combined Enrichr score calculated within GSEApy. **(d)** Circos plot visualisation of selected significant (adjusted p-value <0.05) predicted interactions between TMLM and co-localising neuronal cells in prenatal skin. Arrows represent directionality of interactions (ligand to receptor); connections are coloured by sender cell type with width proportional to the CellphoneDB mean value for each ligand-receptor pair. **(e)** Heatmap of normalised (z-score) mean expression of angiogenesis gene modules in prenatal skin macrophages. CD7hiP: CD7^high^ progenitors, CD7loP: CD7^low^ progenitors, ErP: erythroid progenitors, GMP: granulocyte-monocyte progenitors, HSPC: haematopoietic stem and progenitor cells, ILC: innate lymphoid cells, LC: Langerhans cells, Mac1-4: macrophages 1-4, MkP: megakaryocte progenitors, TML macrophage: *TREM2^+^* microglia-like macrophage, YSMP: yolk-sac derived myeloid-biased progenitors.

**Extended Data Fig. 8.**
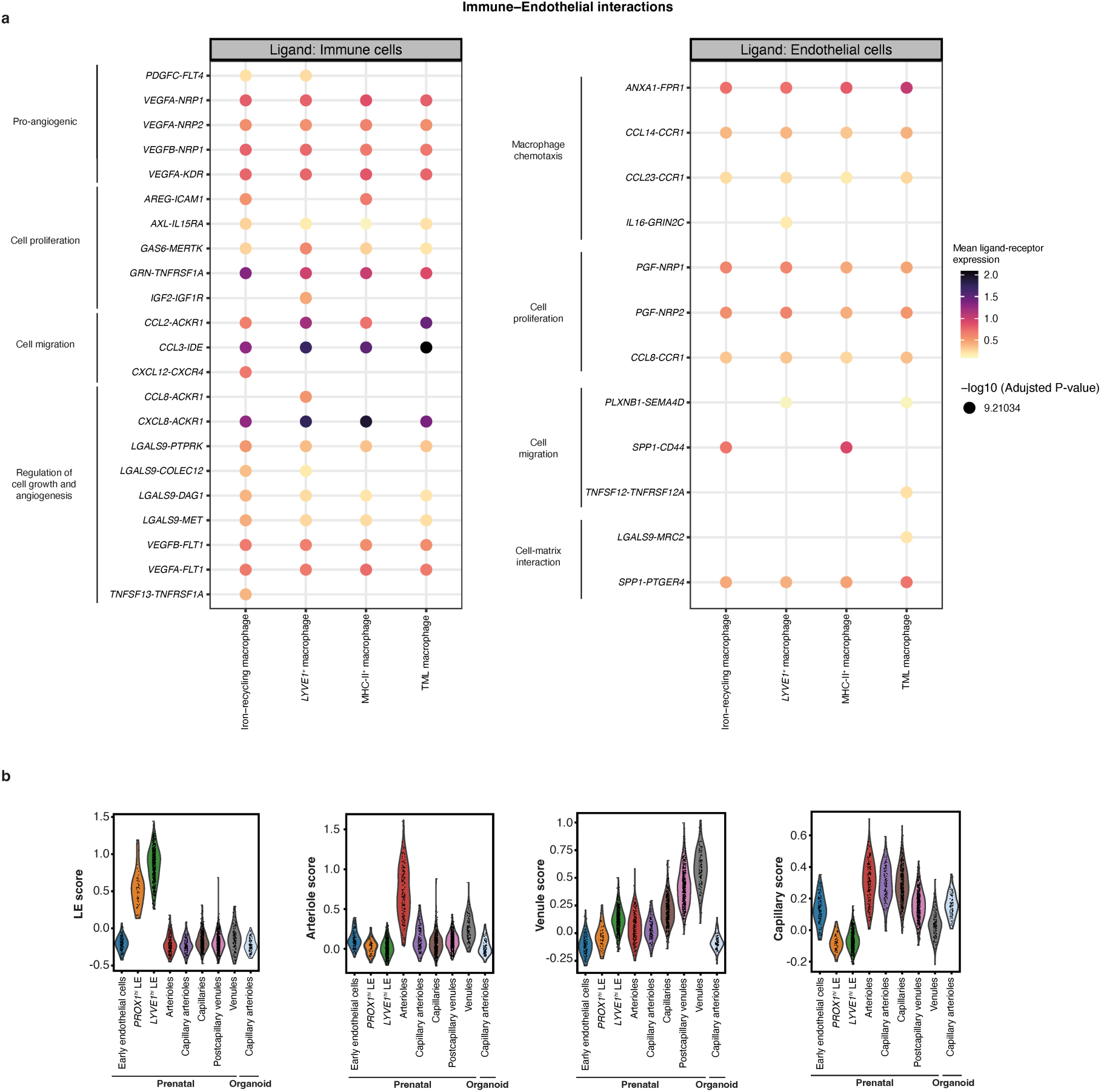
Endothelial cell heterogeneity and interactions with macrophages. **(a)** Dot plot visualisation of selected significant (adjusted p-value<0.05) CellphoneDB-predicted interactions between macrophage subsets and co-localising vascular endothelial cells in prenatal skin, grouped by function. Right: Ligand (first gene in each gene pair) is expressed by macrophages; Left: Ligand (first gene in each gene pair) is expressed by endothelial cells. Dot colour represents the mean expression values of each ligand-receptor pair for the corresponding cell pairs, dot size represents -log_10_(adjusted p-value). **(b)** Violin plots of gene module scores in prenatal skin and skin organoid endothelial cells. Scores were derived from marker genes for the different endothelial cell groups.

**Extended Data Fig. 9.**
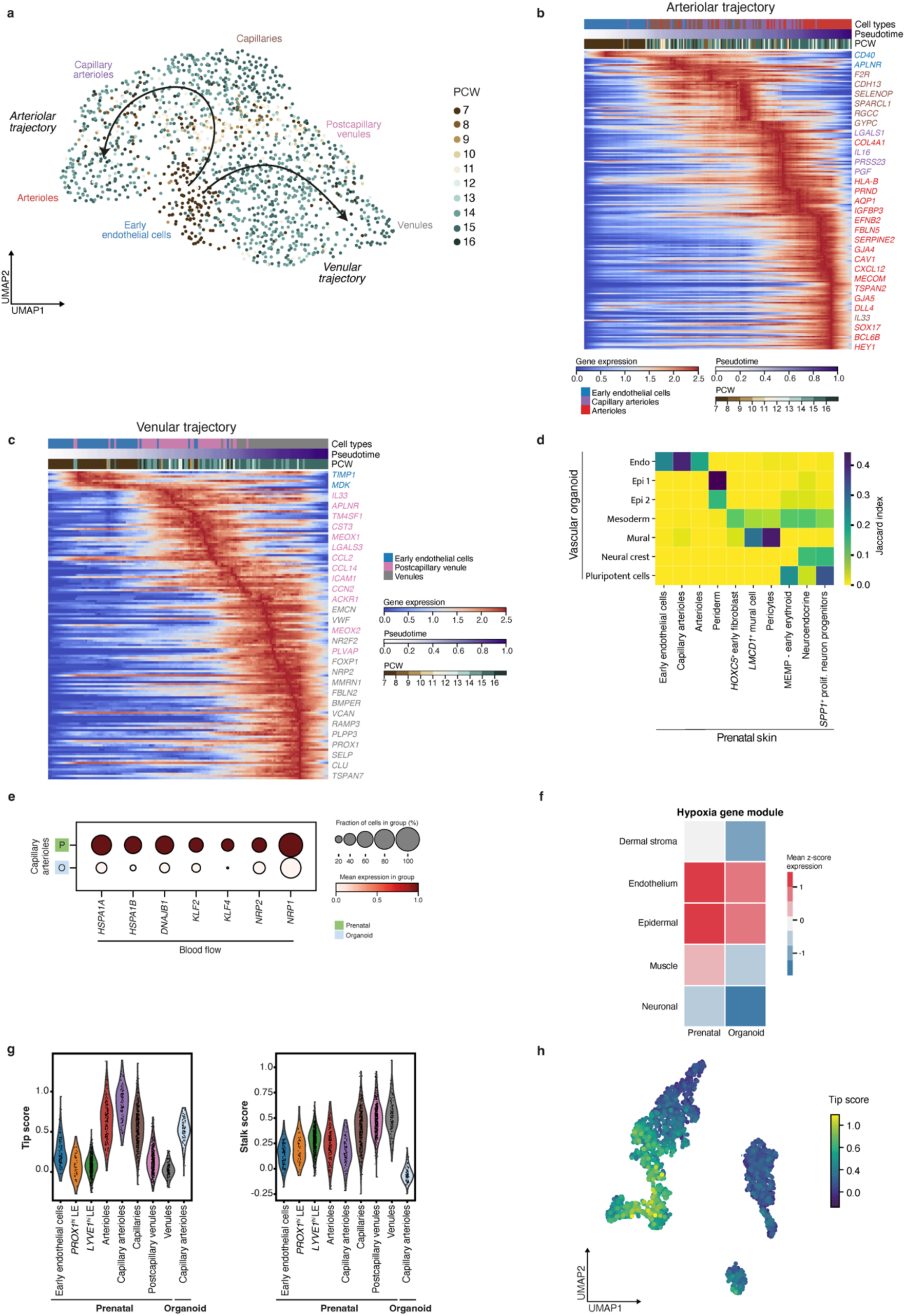
Factors driving angiogenesis and endothelial cell differentiation. **(a)** Inferred pseudotime trajectory of prenatal skin endothelial cell states coloured by gestational age (PCW). **(b)** Heat map showing differentially expressed genes across pseudotime along the ‘arteriolar’ differentiation trajectory. **(c)** Heat map showing differentially expressed genes across pseudotime along the ‘venular’ differentiation trajectory. **(d)** Heatmap showing the correspondence (measured by Jaccard index) between prenatal skin (x-axis) and blood vessel organoid cell states^128^ (y-axis) for a logistic regression model trained on prenatal skin data. The top 10 predicted prenatal cell states were retained for visualisation. **(e)** Dot plot showing variance-scaled, mean expression (dot colour) and percent of expressing cells (dot size) of blood flow-related genes in prenatal skin and skin organoid capillary arteriole cells. **(f)** Heatmap of normalised (z-score) mean expression of hypoxia gene module in prenatal skin and corresponding cell categories in skin organoid. **(g)** Violin plots of ‘Tip’ and ‘Stalk’ cell module scores in prenatal skin and skin organoid endothelial cells. **(h)** UMAP visualisation of the ‘Tip’ cell module score in prenatal skin and skin organoid endothelial cells.

**Extended Data Fig. 10.**
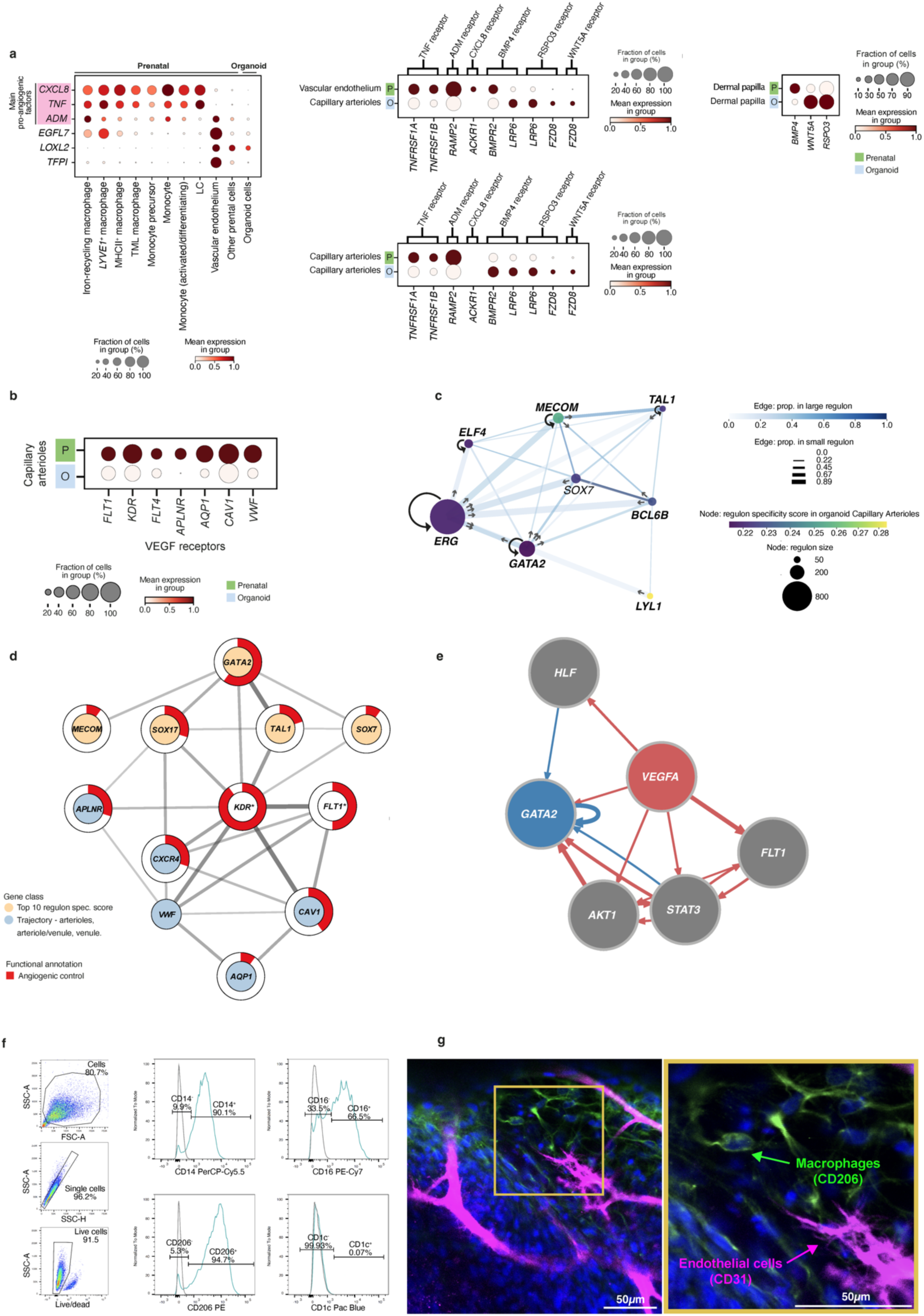
Macrophages support prenatal skin and skin organoid angiogenesis. **(a)** Dot plot showing variance-scaled, mean expression (dot colour) and percent of expressing cells (dot size) of pro- and anti-angiogenic factors and of corresponding receptors in prenatal skin and skin organoid endothelial cells. Genes encoding the main pro-angiogenic factors secreted by macrophages in prenatal skin are highlighted. **(b)** Dot plot showing variance-scaled, mean expression (dot colour) and percent of expressing cells (dot size) of genes (vascular endothelial growth factor receptor and endothelial differentiation genes) in prenatal skin and skin organoid capillary arteriole cells. **(c)** Gene regulation network for five regulons with high specificity score in prenatal skin and/or skin organoid capillary arterioles. Arrows indicate the direction of regulation from transcription factor to target gene. Edges show the proportion of genes shared by two regulons (colour for proportion in the larger regulon and thickness for proportion in the smaller regulon). **(d)** Gene network for five regulons with high specificity score in prenatal skin and/or skin organoid capillary arterioles (*GATA2*, *MECOM*, *SOX17*, *TAL1*, *SOX7*), and selected *GATA2* target genes. The proportion of red in the ring around nodes indicates the proportion of gene ontology terms associated with angiogenesis in the gene set enrichment analysis performed with genes in the network. **(e)** Tree diagram showing network of interactions (NicheNet) linking the ligand *VEGFA* (red) to *GATA2* as target gene (blue) through identified signalling mediators and transcriptional regulators (grey). Edges representing signalling interactions are coloured red and gene regulatory interactions in blue; edge thickness is proportional to the weight of the represented interaction. **(f)** Gating strategy used on iPS-derived macrophages before co-culture (n=1 batch, day 38 of differentiation) to isolate single live cells, analyse expression of macrophage markers (CD14, CD16, CD206) and exclude dendritic cells (CD1c). **(g)** Brightfield images of macrophage and endothelial cell co-culture at 16 hours and 48 hours of the angiogenesis assay suggesting interaction between the two cell types.

